# Structurally complex osteosarcoma genomes exhibit limited heterogeneity within individual tumors and across evolutionary time

**DOI:** 10.1101/2021.08.30.458268

**Authors:** Sanjana Rajan, Simone Zaccaria, Matthew V. Cannon, Maren Cam, Amy C. Gross, Benjamin J. Raphael, Ryan D. Roberts

## Abstract

Osteosarcoma is an aggressive malignancy characterized by high genomic complexity. Identification of few recurrent mutations in protein coding genes suggests that somatic copy-number aberrations (SCNAs) are the genetic drivers of disease. Models around genomic instability conflict - it is unclear if osteosarcomas result from pervasive ongoing clonal evolution with continuous optimization of the fitness landscape or an early catastrophic event followed by stable maintenance of an abnormal genome. We address this question by investigating SCNAs in >12,000 tumor cells obtained from human osteosarcomas using single cell DNA sequencing, with a degree of precision and accuracy not possible when inferring single cell states using bulk sequencing. Using the CHISEL algorithm, we inferred allele- and haplotype-specific SCNAs from this whole-genome single cell DNA sequencing data. Surprisingly, despite extensive structural complexity, these tumors exhibit a high degree of cell-cell homogeneity with little sub-clonal diversification. Longitudinal analysis of patient samples obtained at distant therapeutic time points (diagnosis, relapse) demonstrated remarkable conservation of SCNA profiles over tumor evolution. Phylogenetic analysis suggests that the majority of SCNAs were acquired early in the oncogenic process, with relatively few structure-altering events arising in response to therapy or during adaptation to growth in metastatic tissues. These data further support the emerging hypothesis that early catastrophic events, rather than sustained genomic instability, give rise to structural complexity, which is then preserved over long periods of tumor developmental time.

**Significance Statement:** Chromosomally complex tumors are often described as genomically unstable. However, determining whether complexity arises from remote time-limited events that give rise to structural alterations or a progressive accumulation of structural events in persistently unstable tumors has implications for diagnosis, biomarker assessment, mechanisms of treatment resistance, and represents a conceptual advance in our understanding of intra-tumoral heterogeneity and tumor evolution.

## Introduction

Osteosarcoma is the most common primary bone tumor affecting children and adolescents^1^. Nearly always high grade and aggressive, this disease exhibits extensive structural variation (SV) that results in a characteristically chaotic genome^2–4^. With few recurrent point mutations in protein coding regions, osteosarcoma genomes often exhibit widespread structural complexity, giving rise to associated somatic copy-number aberrations (SCNAs), a likely genomic driver of malignant transformation^5^. Indeed, osteosarcoma is the prototype tumor whose study led to the discovery of chromothripsis^6,7^, a mutational process that causes the shattering of chromosomes leading to localized genomic rearrangements causing extreme chromosomal complexity^8^. However, genomic complexity in osteosarcoma often goes beyond alterations caused by the canonical processes associated with chromothripsis^9,10^. Many have reasonably interpreted chromosomal complexity to be evidence of sustained chromosomal instability (CIN), often with supporting evidence from other cancer types^11–14^. Indeed, cancer sequencing studies have identified the presence of extensive SCNAs as a marker for CIN^13^.

Two distinct models have been proposed to explain the evolution of chromosomal structure and copy numbers in cancer genomes. One model suggests that underlying genomic instability gives rise to populations of cells with diverse phenotypic variations and that ongoing selection of advantageous phenotypes drives evolution and adaptation^15,16^. A somewhat competing model argues that discrete periods of genomic instability, isolated in tumor developmental time, give rise to extreme chromosomal complexity driven by a small number of impactful catastrophic events^17,18^. In osteosarcoma, investigators have put forward data that would seem to support both models. For instance, several groups have used single cell RNA sequencing experiments, which reveal a high degree of transcriptional heterogeneity, to infer a high degree of copy number heterogeneity within osteosarcoma tumors^19,20^, an observation which would support a malignant process driven by ongoing instability and gradual evolution. However, others have shown that SCNA profiles differ little when comparing primary to metastatic or diagnostic to relapse samples^5,21,22^, which would suggest that ongoing mechanisms of malignancy do not create an environment of chromosomal instability. Overall, it remains unclear whether the structurally complex genomes characteristic of osteosarcoma emerge from continuous cycles of diversification and fitness optimization within a context of ongoing instability and significant intra-tumoral chromosomal heterogeneity or from an early catastrophic event that gave rise to widespread structural changes that are then maintained over long periods of tumor development, with evidence from the literature supporting both potential mechanisms.

One challenge in addressing this question comes from challenges in data interpretation and deconvolution, as the existing studies describing copy number clonality and evolution have inferred cell-specific copy number states from bulk tumor sequencing, often from a single time point^23^. However, investigating ongoing clonal evolution from bulk sequencing data remains particularly challenging, as each bulk tumor sample is an unknown mixture of millions of normal and cancer cells^24–27^. The emergence of single cell genomic DNA sequencing technologies now permits scalable and unbiased whole-genome single cell DNA sequencing of thousands of individual cells in parallel^24,28^, providing an ideal framework for analyzing intra-tumor genomic heterogeneity and SCNA evolution. Complementing these technical developments, recent computational advances – most notably the CHISEL algorithm^27^ – enable highly accurate ploidy estimates and the inference of allele- and haplotype-specific SCNAs in individual cells and sub-populations of cells from low coverage single cell DNA sequencing. This allows cell-by-cell assessment of intra-tumoral SCNA heterogeneity, identification of allele-specific alterations and reconstruction of the evolutionary history of a tumor from thousands of individual cancer cells obtained at a single or multiple time points during tumor progression.

Here, we leverage these approaches to determine whether the widespread SCNAs in osteosarcoma result from ongoing genomic instability, providing a mechanism for tumor growth and evolution. Using expanded patient tissue samples, our studies revealed widespread aneuploidy and SCNAs in 12,019 osteosarcoma cells from ten tumor samples. Using this approach, we found negligible intra-tumor genomic heterogeneity, with remarkably conserved SCNA profiles when comparing either the individual cells within a tumor or tumors collected at different therapeutic time points from the same patient. These findings suggest that the widespread patterns of genomic SVs in osteosarcoma are likely acquired early in tumorigenesis, and the resulting patterns of SVs and SCNAs can be preserved within an individual tumor, across treatment time and through the metastatic bottleneck.

## Results

### Individual cells within osteosarcoma tumors exhibit extensive SCNAs, but a high degree of homogeneity

Single cell DNA sequencing was performed on 12,019 tumor cells from expanded patient tissue samples. These nine patient tissues were obtained from diagnostic biopsies of localized primary tumors (*n* = 3), from post-chemotherapy resection procedures (*n* = 2), or from relapsed metastatic lung lesions (*n* = 4), representing a spectrum of disease progression (Supplemental **Table S1**, Supplemental **Table S2**). Apart from OS-17, a well-established model of metastatic osteosarcoma^29^, all patient tissues were expanded for a single passage in mice as either subcutaneous flank or orthotopic bone tumors to obtain fresh tissue to perform single cell DNA sequencing (300-2,500 single cells per sample; Supplemental **Figure S1A**). Previous studies have shown that this procedure yields samples with a high degree of fidelity relative to the diagnostic specimens, especially in early passages, an observation that we also validated in our own samples^30^. We then used CHISEL^27^ to identify allele- and haplotype-specific SCNAs from the sequencing data.

Consistent with previous reports^6,31^, sequencing showed a high degree of aneuploidy and extensive SCNAs across the entire osteosarcoma genome (Figure 1). If driven by a process of chromosomal instability and ongoing/continuous clonal evolution, we would expect to observe multiple subclones with distinct complements of SCNAs within each same tumor, such as has been shown in recent single cell studies of other cancer types^24,27,32^. However, in each of the ten samples investigated, we identified one dominant clone that comprised nearly all cells within each sample, with many samples composed entirely of a single clone (Supplemental **Figure S1B**). To ensure that our results were not an artifact caused by the algorithm or the selected thresholds for noise control, we confirmed that the cells discarded as poor quality/noisy by CHISEL bear SCNAs similar to the dominant clones identified in each sample – thus no rare clones with distinct copy number profiles were discarded inappropriately (Supplemental **Figure S2**). Interestingly, we found that a substantial fraction of the overall copy-number changes involved allele-specific SCNAs, including copy-neutral LOHs (i.e., allele-specific copy numbers {2, 0}) that would have been missed by previous analyses of total copy numbers.

**Figure 1.**
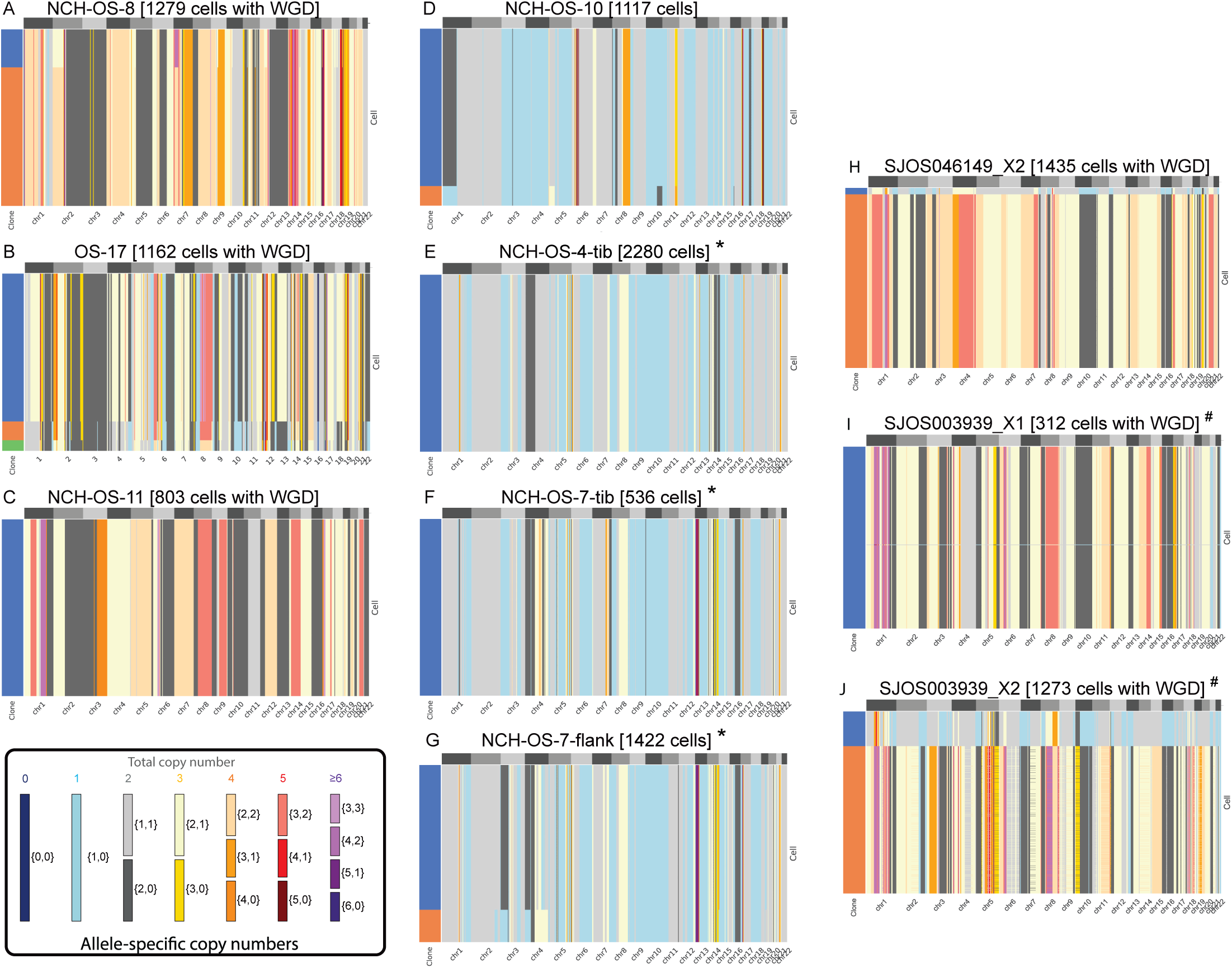
Extensive genomic complexity in ten expanded osteosarcoma patient tissue samples using single cell DNA sequencing. Allele-specific copy numbers (heatmap colors) are inferred by using the CHISEL algorithm^27^ from each of ten datasets including 300-2300 single cancer cells from osteosarcoma tumors. In each dataset, cancer cells are grouped into clones (colors in leftmost column) by CHISEL based on the inferred allele-specific copy numbers. Corrected allele-specific copy-numbers are correspondingly obtained by consensus. Note that cells classified as noisy by CHISEL have been excluded. ‘*’ and ‘#’ represent samples obtained from the same patient.

Genome-wide ploidy of single cells showed high variability across samples, ranging from 1.5 to 4, demonstrating a high degree of aneuploidy (Supplemental **Figure S3**). Consistent with the high levels of aneuploidy, we identified the presence of whole-genome doubling (WGD, a phenomenon identified with much greater precision in the single cell data) across nearly all cancer cells of six tumors (NCH-OS-8, OS-17, NCH-OS-11, SJOS046149_X2, SJOS003939_X2 and SJOS003939_X1; **Figure 1 A-C, H-J**). One tumor (SJOS003939_X2) shows two subclones that appear to be undergoing whole genome duplication, with one subclone exhibiting a SCNA pattern that is almost exactly double that of the other, across the genome.

To further assess tumor stability, we used paired datasets from patients collected at time points along tumor progression. We observed that whole-genome copy-number profiles were highly consistent within each patient. The first set includes NCH-OS-4, which was obtained shortly after diagnosis at the time of resection (after two rounds of neoadjuvant MAP chemotherapy), and NCH-OS-7, which was obtained at the time of relapse the following year. Comparing genomic windows where at least one sample had a SCNA in the primary clone, 77-78% of genomic windows had identical copy number assignments in both samples, despite variation in tumor purity (**Supplemental Figure S4**). This contrasts with between 1% and 35% concordance for non-related samples. The correlation between related samples may be even higher, given inaccuracies expected from low-coverage single cell SCNA detection.

The second set of paired primary and metastatic lesions (SJOS003939_X1, SJOS003939_X2) also showed SCNA profiles that were highly similar (58% of SCNAs identical, **Supplemental Figure S4**), suggesting a high degree of conservation of genomic aberration profiles over therapeutic time. Overall, we observed a very high degree of homogeneity within cancer cells sequenced from the same tumor. Even in tumors where small proportions of cells (5-20%) are classified as part of small subclones, these sub-clonal cells are only distinguished by modest SCNAs differences within a few chromosomes. The exception to this general observation arose in SJ0S003939_X2, a second xenograft from a patient with a germline TP53, raising suspicion for a second malignancy (rather than a relapse). Thus, despite the high levels of aneuploidy and massive SCNAs identified in all ten samples, these osteosarcoma cells demonstrated very modest levels of intra-tumor heterogeneity and variation across therapeutic time.

### Osteosarcoma cells harbor extensive SCNAs that mostly correspond to deletions

The occurrence of WGD events correlates with high levels of aneuploidy and higher frequency of SCNAs^33^. Recent reports have identified that WGDs serve as a compensatory mechanism for cells to mitigate the effects of deletions^34^. We investigated cell ploidy and fraction of genome affected by SCNAs (aberrant), amplifications, deletions, and sub-clonal CNAs between tumors affected by WGDs (NCH-OS-8, OS-17, NCH-OS-11, SJOS046149_X2, SJOS003939_X2 and SJOS003939_X1) and tumors not affected by WGDs (NCH-OS-10, NCH-OS-4 and NCH-OS-7). Osteosarcoma cells in all analyzed tumors demonstrate extensive SCNAs, affecting more than half of the genome in every tumor cell. We found that the fraction of genome affected by SCNAs ranged from 50-70% on average (**Figure 2A**, Supplemental **Figure S5A**). This result might not be surprising for tumors affected by WGDs, however, we observed that tumors not affected by WGD had a high fraction of aberrant genome as well (higher than 50% on average; **Figure 2A**). This aberrant fraction is substantially higher than has been reported for other cancer types^33^.

**Figure 2.**
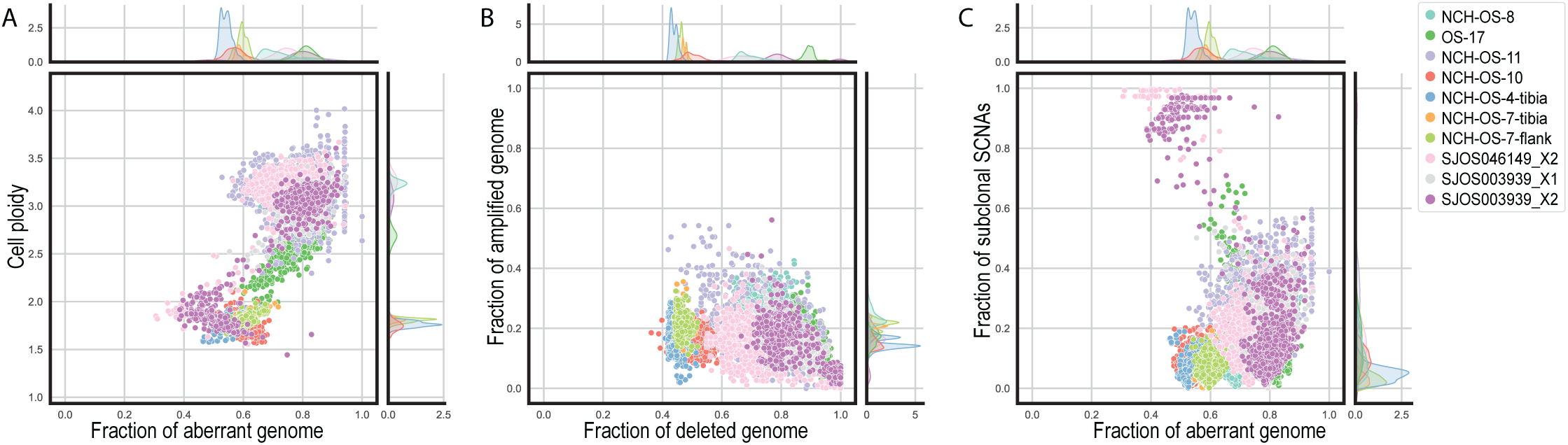
Osteosarcoma cancer cells exhibit extensive genetic alterations, especially deletions, but a relatively low level of heterogeneity. (A) Ploidy (y-axis) and fraction of aberrant genome (x-axis) of every cell (point) across the ten analyzed datasets (colors). The kernel density of the marginal distributions of each value is reported accordingly in every plot. (B) Fraction of genome affected by deletions (x-axis) vs. fraction of genome affected by amplifications (y-axis) of every cell (point) across the ten analyzed datasets (colors). (C) Fraction of aberrant genome (x-axis) and fraction of sub-clonal SCNAs (i.e. fraction of the genome with SCNAs different than the most common clone for the same region across all cells in the same dataset, y-axis) of every cell (point) across the ten analyzed datasets (colors).

We observed a clear enrichment of deletions among the identified SCNAs across all cancer cells. The fraction of genome affected by amplifications is 0-40% on average in every tumor, while the fraction of the genome affected by deletions is 40-100% on average across all cancer cells in every tumor (**Figure 2B**). This result is not particularly surprising for tumors with WGD events and is consistent with a recent study of Lopez et al.^34^ that demonstrated a similar correlation in non-small-cell lung cancer patients. However, in the osteosarcoma tumors analyzed in this study, we found that cancer cells in non-WGD tumors are similarly affected by a high fraction of deletions (**Figure 2B**). Importantly, we observed that >80% of the cells in all but two of our samples harbored LOH events at the TP53 locus (in-line with frequency previously reported^3^) (**Supplemental Figure S5**). This substantiates the correlation between LOH of TP53 and high levels of genomic instability (including the occurrence of WGDs) reported in recent studies^13,34,35^, and suggests that these events might have a critical role in the maintenance of a highly aberrant genomic state. Notably, CHISEL identified 50% of the samples to harbor copy-neutral LOH alterations at the TP53 locus that would be missed by total copy number analyses (**Supplemental Figure S5**).

We found it interesting that sub-clonal SCNAs that likely occurred late in the evolutionary process (present only in subpopulations of cancer cells) are relatively rare across all analyzed osteosarcoma tumors, irrespective of WGD status (with a frequency of 0-20% in most cancer cells; **Figure 2C**, **Supplemental Figure S5**). Note the only exceptions to this observation correspond to cells in NCH-OS-11, a sample with overall higher noise and variance, and a subpopulation of cells in two other tumors (SJOS046149_X2 and SJOS003939_X2) that appear to be cells that have not undergone WGD (**Supplemental Figure S5B**). Indeed, the average fraction of SCNAs in SJOS046149_X2, SJOS003939_X2 is lower than 20%. Overall, we observed that osteosarcoma cells investigated in these ten samples, whether passaged in cell culture over a few generations (OS-17), treatment naïve or exposed to extensive chemotherapy, bear high levels of aneuploidy marked with extensive deletions and negligible sub-clonal diversification, irrespective of WGD status.

### Longitudinal single cell sequencing shows modest evolution of SCNA from diagnosis to relapse

Increased aneuploidy has previously been associated with chromosomal instability (CIN) and accelerated tumor evolution^13,36^, though some have suggested that this observation specifically applies to tumors that exhibit not only high levels of SCNA, but also high levels of sub-clonal SCNA^37^. To assess the degree of structural instability exhibited by these tumors, we examined a pair of samples, NCH-OS-4 and NH-OS-7, collected at diagnosis and at relapse respectively, from the same patient to determine whether SCNAs remained stable over therapeutic time or showed signs of significant instability/evolution. This included an expansion in both the flank and orthotopic tibia locations to determine whether these environments drove a niche-specific expansion of a selected clone. Results suggest that expansion in mouse did not lead to evolutionary disequilibrium.

We used CHISEL to jointly analyze 4,238 cells from these paired tumor samples and to infer corresponding allele- and haplotype-specific SCNAs (**Figure 3A**). Based on existing evolutionary models for SCNAs, we reconstructed a phylogenetic tree that describes the evolutionary history of the different tumor clones identified in these tumors (**Figure 3B**). The result from this phylogenetic analysis confirmed our findings in two ways. First, we found that the evolutionary ordering of the different clones in the phylogenetic tree is concordant with the longitudinal ordering of the corresponding samples (**Figure 3B**): the tumor clones identified in the early sample (NCH-OS-4) correspond to ancestors of all the other tumor clones identified in later samples (NCH-OS-7-tib and NCH-OS-7-flank). Second, we observed that the vast majority of SCNAs accumulated during tumor evolution are truncal, indicating that these aberrations are accumulated early during tumor evolution and shared across all the extant cancer cells (**Figure 3B**). In fact, only three significant events distinguish the most common ancestor of all cells from this patient (identified in NCH-OS-4) from the cells within the relapse lesion: gain of chromosome 14, gain of chromosome 16q (resulting in copy-neutral LOH), and deletion of one allele of chromosome 18 (resulting in LOH). Note that we cannot be certain of when these clones arose. It is possible these changes occurred early in tumor formation and were present in the primary tumor but were not present in the biopsied sample and so we must exercise caution when assessing if there is ongoing low-level chromosomal instability.

**Figure 3.**
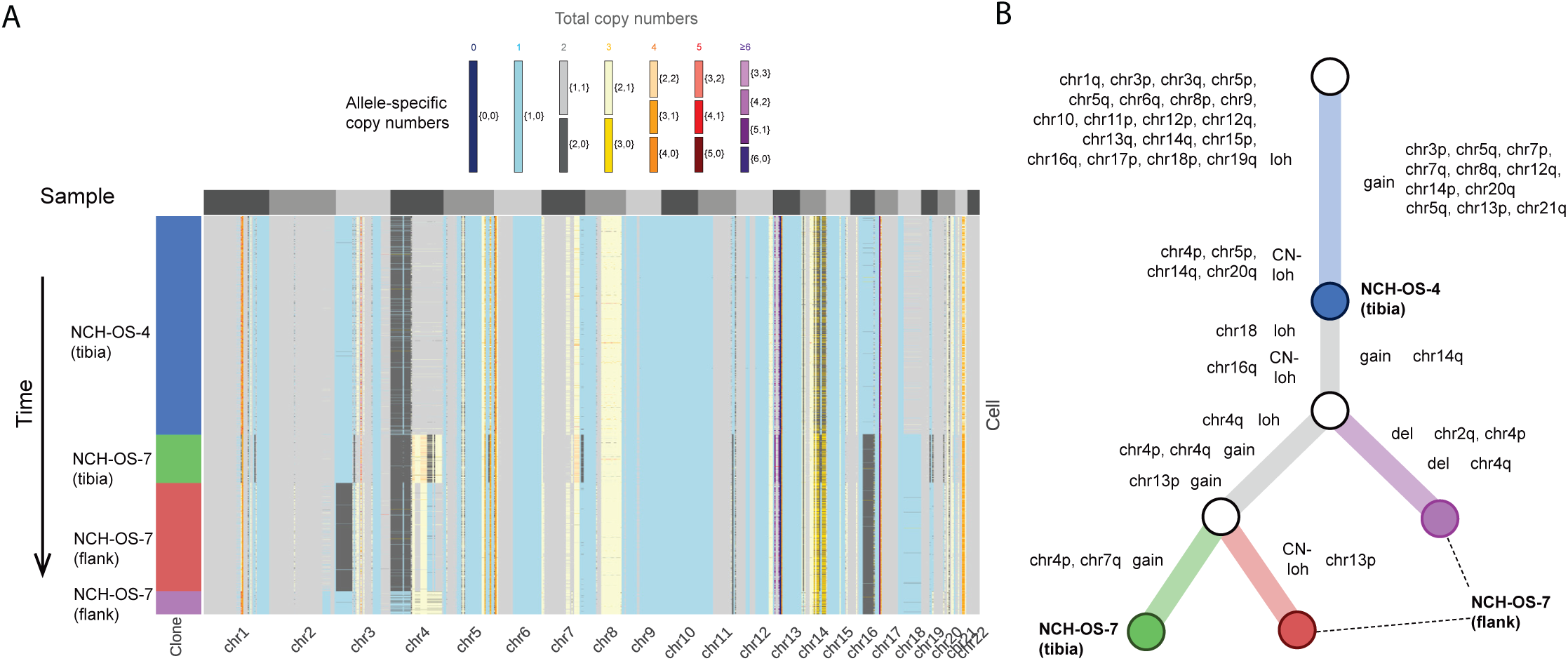
Phylogenetic reconstruction of tumor evolution is consistent with longitudinal ordering of matched tumor samples and reveals conservation of SCNA profiles. (A) Allele-specific copy numbers (heatmap colors) across all autosomes (columns) have been inferred by CHISEL jointly across 4238 cells (rows) in 3 tumor samples from the same patient: 1 pre-treatment sample (NCH-OS-4 tibia) and two post-treatment samples (NCH-OS-7 tibia and NCH-OS-7 flank). CHISEL groups cells into 4 distinct clones (blue, green, red, and purple) characterized by different complements of SCNAs. (B) Phylogenetic tree describes the evolution in terms of SCNAs for the four identified tumor clones. The tree is rooted in normal diploid clone (white root) and is characterized by two unobserved ancestors (white internal nodes). Edges are labelled with the corresponding copy-number events that occurred and transformed the copy-number profile of the parent into the profile of the progeny. The four tumor clones (blue, green, red, and purple) are labelled according to the sample in which they were identified.

To further assess the effects that environmental stressors might play on the creation and/or emergence of sub-dominant clones, which could be masked due to extreme rarity, we expanded samples from the same tumor within two different microenvironments in mice. Consistent with the diagnosis-relapse sample comparison, clones identified within tumors grown orthotopically within the tibia (NCH-OS-7-tib) or within subcutaneous flank tissues (NCH-OS-7-flank) are highly concordant (78% of genomic windows called with identical SCNA values) and distinguished by few focal SCNAs (primarily single copy changes). These changes could be either be variance in SCNA calling from the sequencing data, stochastic differences caused by the presence of sub-clones within the original tumor sample that was bisected and implanted or biologically relevant. Without targeted studies it is not possible to confidently define the biological role of these focal changes, if any. A third comparison of tumors separated in time and space was possible using another paired set of primary and metastatic lesions (SJOS003939_X1, SJOS003939_X2), which also demonstrated negligible sub-clonal diversification (**Supplemental Figure S7**). Indeed, each sample was dominated by one major clone, which exhibited only subtle differences from the paired sample. While these results do not suggest the absence of SCNA changes over the course of tumor evolution, they do suggest a level of stability quite similar to genomically simple tumors and that the mechanisms giving rise to these limited focal changes are different from those that gave rise to widespread genomic complexity.

To further explore temporal and spatial consistency of patient tumor samples, we combined whole genome sequencing data obtained from paired osteosarcoma samples within the St. Jude database^38–40^ with our own whole genome sequencing and performed SCNA analysis. This combined data yielded between two and six tumor samples for each patient, in addition to a germline reference sample. In most cases, all samples taken from a single patient at different timepoints were highly similar and clustered together (**Figure 4** and **Supplemental Figure S8A-K**). There were, however, five samples that had more than one distinct clone in separately collected samples which reduced the overall average. In these instances, the average correlation between clonal populations within a patient was only 0.28, which was close to the correlation we observed between samples taken from different patients (mean Pearson cor = 0.18). Deeper exploration of these samples revealed germline TP53 mutations in some patients (shown with a red asterisk in **Figure 4**), suggesting an underlying cancer predisposition and a likelihood that these are tumors arising from distinct oncogenic events. The correlation within a clone was very high (mean Pearson cor = 0.67), despite the noise created by the sparse coverage sequencing inherent to this method. Xenograft-derived samples did not cluster separately from samples derived directly from patients (**Figure 4**), except for two samples from SJOS030645 which formed a distinct cluster. The xenograft samples had a high correlation with non-xenograft samples from the same patient (mean Pearson cor = 0.70), suggesting that the SCNA patterns in these samples were not dramatically altered by clonal selection within the mouse. Determination of SCNA-based clonal composition and tumor purity was performed using the HATCHet algorithm^41^, providing additional context for interpretation of results. HATCHet results show a very high degree of SCNA-based clonal conservation from one clinical timepoint to the next.

**Figure 4.**
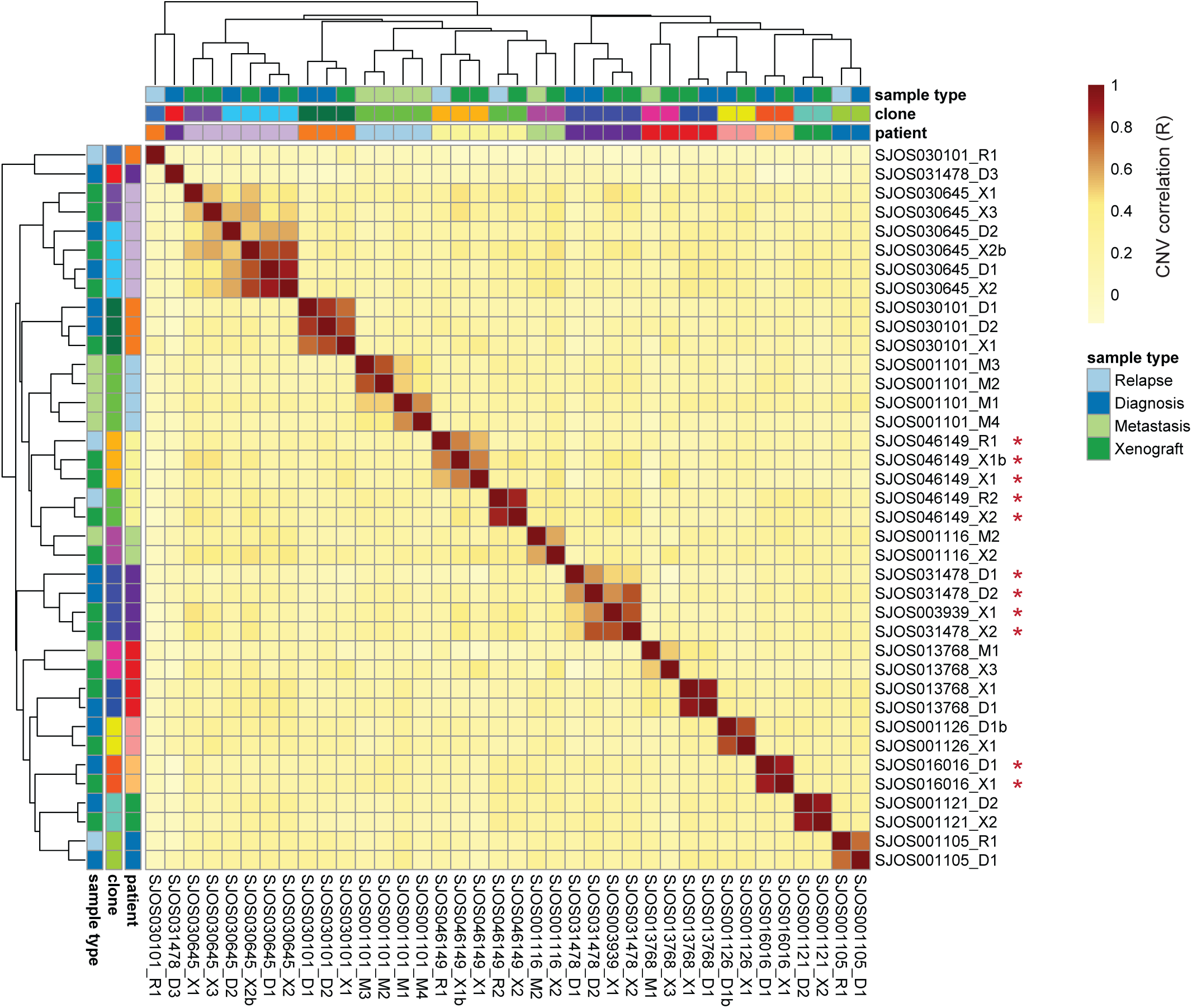
CNA correlation between osteosarcoma samples. Pearson R values denoting correlation of binned copy numbers between samples. Colors on x and y-axes indicate each sample’s patient of origin and type as well as the clones defined from the correlation analysis. Red asterisks denote samples from patients with germline TP53 mutations. Note that SJOS003939_X1 is from the same patient as SJOS031478_* samples.

## Discussion

Osteosarcoma is one of several malignancies typified by chaotic genomic landscapes dominated by structural variation and corresponding copy number alterations^3^. Chromosomal complexity in osteosarcoma and other cancers with complex karyotypes has often been assumed to represent underlying genomic instability, suggesting that these tumors gradually accumulate structural changes that lead to increasing complexity, with continual selection of ever more aggressive clones driving malignant progression. This concept was supported by previous reports demonstrating that, in some patients, spatially separated tumor samples exhibit slightly divergent SNP and SCNA patterns ^42,43^. These studies also noted that, while there is heterogeneity between samples, there seem to be clones that are shared across multiple metastatic foci. By nature, these studies understandably focused on identifying SNP and copy number differences contained within distinct lesions in these highly aggressive tumors, with the largest sample sets collected at autopsy. By utilizing single cell DNA sequencing, we have been able to investigate intra-tumor genomic heterogeneity and tumor evolution in concrete ways that were previously possible only by estimation and inference using bulk sequencing methods^44–47^. This allowed us to ask different questions related to the stability of these complex genomes within a tumor sample. We were surprised to find that cells within a tumor demonstrated surprisingly little cell-to-cell variability in SCNA profiles - results that, at first, seemed discordant with previous reports of intra-tumoral genomic heterogeneity in osteosarcoma^19^.

Analyzing longitudinal sets of paired samples, we showed that these particular osteosarcoma tumors maintained relatively stable SCNA profiles from diagnosis to relapse, primary tumor to metastasis, and during growth in two distinct environments. Our phylogenetic analysis suggested that the most recent common ancestor of these related samples harbored almost all of the observed SCNAs, suggesting that most of the genomic aberrations arose early in the tumorigenic process within these patients, followed by a long period of stable clonal expansion (clonal stasis)^18^. Our analysis of bulk whole genome sequence data from St. Jude supports this observation and highlights an important observation. Where we observed multiple clones in our single cell data, each clone was homogeneous in its SCNA patterns across cells within the clone, but highly distinct from other clones (**Figure 1**). We observed the same phenomenon in the bulk St. Jude data where a single clone detected in multiple temporally or anatomically separated samples had highly conserved SCNAs while distinct clones were highly divergent (Figure 4). This suggests that each of these clones either derive from a very early event that produced multiple distinct clones, or independent tumorigenesis events.

An inherent limitation of single cell analysis of biopsy samples is that they are not representative of the entire tumor and so the homogeneous cell populations we observe could, in part, derive from the small sample size involved. However, our data include independent data from multiple biopsies that showed similar clonal patterns. Also, our analysis of the St. Jude samples includes multiple independent biopsies from patients and demonstrates the same pattern of SCNA conservation across samplessingle cell. A potential unexpected advantage of the small sample size inherent to tumor biopsies is that these samples tended to be clonal in nature in our single cell data. Given this, it may be feasible to assume that bulk SCNA results are representative of most cells within the biopsy.

Another limitation of biopsy samples is the potential for normal cell types within the sample to interfere with the evaluation of SCNAs. If too large of a proportion of normal cells are present, estimates of copy number will be less accurate. For instance, Supplemental **Figure S8A** shows sample SJOS031478_D1 which has very low copy number alteration values, suggesting that this sample may have a large proportion of normal cells, making detection of SCNA difficult. Deconvolution can improve, but not fully overcome, this issue^41^. To help compensate for this issue, for **Figure 4** we used correlation between samples instead of comparison of absolute copy numbers. This allows sample SJOS031478_D1 to cluster closely with SJOS031478_D2 in **Figure 4** despite apparent contamination with normal tissue. Samples SJOS031478_D1 and SJOS031478_D3 had very low correlation in their SCNA patterns. It is notable that these samples harbor distinct SCNAs including a deletion of a large portion of chromosome five in SJOS031478_D3 that is absent from SJOS031478_D1 and a large amplification of chromosome eight present in SJOS031478_D1 but absent from SJOS031478_D3 (Supplemental **Figure S8A**) indicating that these are distinct clones. Similar patterns can be seen in Supplemental **Figure S8J** where SJOS046149_R2 and SJOS046149_X2 are distinct from SJOS046149_R1, SJOS046149_X1 and SJOS046149_X1b.

One genomic change that was readily evident within our data was the common occurrence of WGD. Using the CHISEL algorithm^27^, we identified high levels of aneuploidy and extensive genomic aberrations that were dominated by deletions within these osteosarcoma tumors. Consistent with previous reports suggesting WGD as a mechanism to mitigate the effects of widespread deletions^34^, we identified extensive deletions even in tumors that had not undergone duplication. Indeed, some of our samples showed subclones of cells that differed across the genome by almost exactly two-fold, which may represent populations of cells that had undergone duplication (with the duplicated fraction being the dominant clone). These findings support the hypothesis that duplication is a process that produces a more aggressive clone from cells that are first affected by widespread deletion.

To expand the analysis addressing the question of stability beyond our single cell WGS samples, we evaluated SNCAs in bulk WGS data derived from osteosarcoma samples. We investigated paired tumor samples across 14 patients (and included the associated patient-derived xenografts, where available) to determine if SCNA patterns were stable across time. We observed that there were both identical and divergent clones within single patients. Clones were similar with correlations as high as 0.92. In the few patients where relapse specimens contained highly divergent clones (**Supplemental Figure S8A-K**), a deeper analysis revealed germline TP53 mutations in many cases (**Figure 4**). In these patients harboring a genetic predisposition to developing osteosarcoma, it is likely that these genetically distinct lesions represent independent oncogenic events and it is possible that TP53 activity was impaired through alternative means in the other patients.

Historically, studies in osteosarcoma (and other cancers) have equated a high level of SCNA with ongoing genomic instability^31,48^, and some direct evidence has supported this concept^43^. However, several recent studies seem to challenge this conclusion, showing preservation of SCNA profiles in primary vs metastatic and diagnostic vs relapse samples^5,22,42^. Our findings support the hypothesis that mechanisms leading to widespread structural alterations are active early in tumorigenesis but resolve and are followed by long periods of relative stability. These seemingly discordant observations may both be true. First, there may be different paths to chromosomal complexity in different tumors—processes that resolve in some tumors, but do not in others. Indeed, nearly all these publications contain sample sets that seem to support higher and lower levels of chromosomal (in)stability.

Second, tumor cells may experience time-limited periods of relative instability, resulting in the phenomenon of punctuated equilibrium, as has been shown in other cancer types^18^. In a punctuated equilibrium scenario, the timing of the biopsy would change the likelihood of finding more or less SCNA heterogeneity within the tumor using methods like single cell WGS.

To synthesize these concepts, there are several potential models for the emergence of SCNA-defined clones in osteosarcoma (**Figure 5**). A single initiation event giving rise to a single dominant clone followed by highly stable genomic organization (**Figure 5A**) would cause all tumor samples from a patient to have consistent SCNA patterns in both bulk and single cell sequencing. This mechanism, however, would not be supported by previously published data ^42,43^ or the results presented here.

**Figure 5.**
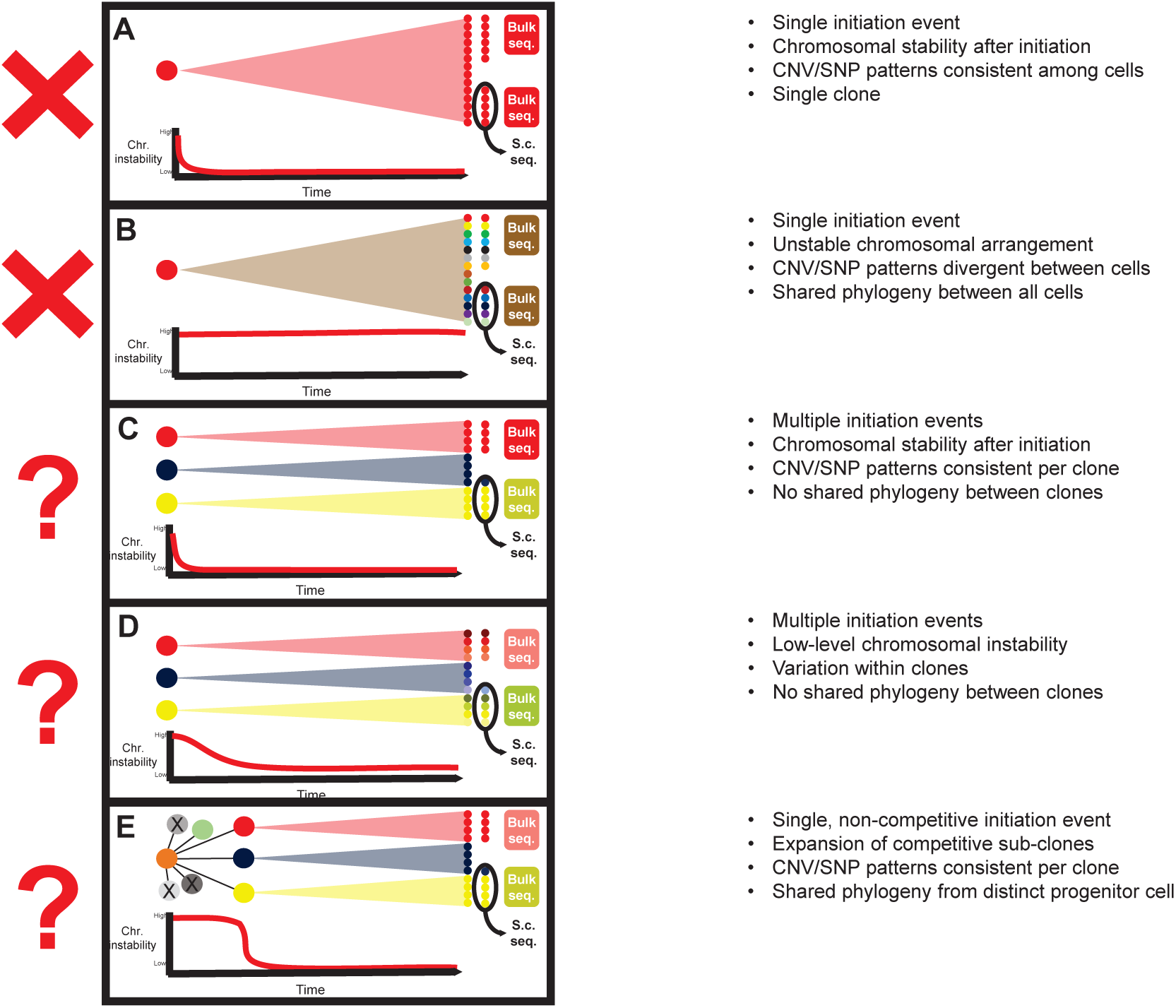
Possible models for temporal SCNA stability. (A) After tumor initiation, if chromosomal instability is low, tumors will have identical SCNA patterns across all cells. (B) High tumor instability would result in tumors with highly heterogeneous SCNA patterns across cells which may not be apparent in bulk sequencing. (C) If there are multiple initiation events with low subsequent genomic instability SCNA patterns will be consistent across clones, which will be apparent by both bulk and single cell sequencing. (D) If there are multiple initiation events with ongoing genomic instability, clones derived from each will be similar but with highly variable SCNA patterns within a clone. This heterogeneity would be apparent by single cell sequencing but not bulk sequencing. (E) If a single initiation event is followed by an initial period of genomic instability, divergent clones could emerge. Patterns of stability within a clone would suggest that chromosomal stability is re-established prior to clonal expansion.

In an alternative model, instability persists from the oncogenic insult forward (**Figure 5B**). This model would produce samples containing multiple divergent subclones, which would be evident in bulk sequencing collected in different loci, though not detectable within a single sample. Single cell analyses would identify several SCNA-defined clones within each sample. This model seems less likely considering our single cell results.

A third outcome could result if there were multiple independent initiation events, producing several competing clones within a tumor, followed by a period of stable chromosomal organization (**Figure 5C**). This mechanism, which is consistent with the punctuated equilibrium hypothesis^18^, could produce multiple samples from a patient with divergent SCNA patterns by bulk sequencing, however, cells within each sample would demonstrate highly similar copy number patterns (assuming the sample does not overlap a boundary between clones). This model agrees with both published observations^42,43^ and the results presented here.

A slight modification of this model would invoke early mechanisms giving rise to multiple competing clones, followed by a period where an independent mechanism causes ongoing low-level chromosomal instability within each founder clone (**Figure 5D**). This mechanism would generate multiple slightly divergent clones within each sample. We see some evidence to support this model, such as the similar, but distinct, clones observed in the NCH-OS-7 samples in Figures 1 and 3. These patterns could also be explained by tissue sampling bias, experimental variance or computational noise caused by the low sequencing depth inherent to single cell data. A larger study would be needed to evaluate this.

A final model, which would also be consistent with both our single cell data and the published record, suggests a single initiation event followed by a period of where daughter cells exhibit chromosomal instability (**Figure 5E**), creating a diversity of competing clones. Eventually, clones emerge that exhibit chromosomal stability and have a competitive advantage. In this scenario, tumor cells within a patient are distinct from clone to clone, but homogeneous within a clone, with a small subset of shared SCNAs that were present in the origin cell and maintained through the subsequent chromosomal instability.

While copy number patterns might be stable after an initial structure-altering event, SNVs arise through completely different mechanisms and likely have different evolutionary dynamics. Previous studies looking at osteosarcoma across therapeutic time show clear sequence related changes dominated by patterns that suggest a cisplatin induced mutational burden ^43^. However, the structural integrity of the chromosomes does not seem to be affected by treatment^42,43^.

A brief (single cycle) expansion of tissue using an animal host proved useful for generating high-quality single cell suspensions of sufficient quantity while maintaining high fidelity to the original patient sample. This approach is not intended to model tumor progression in a murine host, but rather to maximize the data obtained from each of these incredibly valuable samples. Some have expressed concerns that mouse-specific evolution selects for sub clonal populations^49^. However, the mouse-specific evolution that occurs over many passages (such as in the development of a PDX) does not occur when the mouse is used as a vehicle for brief expansion^30^. Our SCNA analysis comparing results of these expanded tissues to bulk sequencing performed directly on the patient samples showed a very high correlation between expanded and primary samples. Therefore, this approach may represent a productive compromise enabling multiple lines of research on tissues with limited availability in rare diseases.

Our findings of clonal stasis in osteosarcoma sheds some light on the complex evolutionary history of this cancer type and could have important implications for tumor evolution, patient diagnosis and treatment of osteosarcoma. However, a much larger sample size of patient tissues is needed to capture the full heterogeneity of osteosarcoma seen in the human disease and describe the prevalence of multiple tumor sub-clones. Somewhat ironically, one may conclude from this data that bulk sequencing methods likely produce an adequate assessment of SCNA profiles and heterogeneity in osteosarcoma, given the lack of heterogeneity found in our analysis. These data likewise suggest that, in a clinical setting, sequencing analyses based on SCNA likely remain valid, even into treatment and relapse, assuming separate samples derive from the same clonal tumor population.

At a biological level, these results support the early catastrophe model as a primary mechanism of osteosarcoma complexity, suggesting that most structural rearrangements occur early in the tumorigenic process. While other rearrangements certainly can occur during malignant progression, subsequent structural events do not appear to be *necessary* for invasion, metastasis, or therapeutic resistance (though they certainly may *contribute* to such processes), nor do they appear to be the same mechanisms that create widespread structural complexity. Ongoing research will continue to inform our understanding of the contributions that initial catastrophic events and ongoing mechanisms of genomic evolution have and how they influence clinical outcomes.

It is important to note that our study is performed in a way that is generally insensitive to other alterations (such as SNVs) as a source of genomic variation, though few recurrent mutations have been identified in osteosarcoma, despite extensive genetic analysis^5,48,50^. If both observations hold true, one must conclude that the acquisition of traits that drive malignant progression arise through epigenetic-based evolutionary processes, which remain poorly understood. Interestingly, we and others have shown that these same osteosarcomas demonstrate a high level of intra-tumor transcriptional heterogeneity^19,51^. This heterogeneity of gene expression in cells that are genomically homogeneous suggests that there may be microenvironmental differences or an underlying epigenetic heterogeneity, which could be a basis for competition and selection of tumor cells.

## Materials and Methods

### Experimental model – Expanded patient tissues and murine studies

#### Expanded patient tissue

Patient samples NCH-OS-4, NCH-OS-7, NCH-OS-8, NCH-OS-10 and NCH-OS-11 were obtained from patients consented under an Institutional Review Board (IRB)-approved protocol IRB11-00478 at Nationwide Children’s Hospital (Human Subject Assurance Number 00002860). Germline whole genome sequencing (WGS) was generated from patient blood collected under IRB approved protocol IRB11-00478. Patient samples SJOS046149_X1, SJOS046149_X2, SJOS003939_X1 and SJOS031478_X2, with matched normal WGS were received from St. Jude’s Children’s Research Hospital through the Childhood Solid Tumor Network^38–40^. The OS-17 PDX was established from tissue obtained in a primary femur biopsy performed at St. Jude’s Children’s Research Hospital in Memphis and was a gift from Peter Houghton^29^.

#### Murine Studies. Flank tumors

Viable tissue fragments from patient tissue were expanded in C.B-17/IcrHsd-Prkdc^scid^ mice as subcutaneous tumors following approved IACUC protocols. These tumors were allowed to grow to 300-600 mm^3^ before harvest. Passage 1 expanded tissue was used for all samples, with the exception of OS-17 (p18).

#### Orthotopic primary tumors

Single cell suspensions of 5×10^5^ cells were injected intratibially in C.B-17/IcrHsd-Prkdc^scid^ mice as per IACUC guidelines. These tumors were harvested once they grew to 800 mm^3^ and prepped for single cell DNA-seq.

### Single cell suspension and DNA library generation

Tumors harvested from mice were processed using the human tumor dissociation kit (Miltenyi Biotec, 130-095-929) with a GentleMacs Octo Dissociator with Heaters (Miltenyi Biotec, 130-096-427). Single cell suspensions in 0.04% BSA-PBS of dissociated tumor tissues were generated and frozen down using the 10X freezing protocol for SCNA. The frozen down single cell suspensions were processed using the Chromium Single Cell DNA Library & Gel Bead Kit (10X genomics #1000040) according to the manufacturer’s protocol with a target capture of 1000-2000 cells. These barcoded single cell DNA libraries were sequenced using the NovaSeq 6000 System using paired sequencing with a 100bp (R1), 8bp (i7) and 100bp (R2) configuration and a sequencing coverage ranging from 0.01X to 0.05X (~0.02X on average) per cell. Germline WGS was performed on NovaSeq SP 2×150BP.

### Single cell SCNA calling using CHISEL

Paired-end reads were processed using the Cell Ranger DNA Pipeline (10X Genomics), obtaining a barcoded BAM file for every considered single cell sequencing dataset. As described previously^27^, the pipeline consists of barcode processing and sequencing-reads alignment to a reference genome, for which we used hg19. We applied CHISEL (v1.0.0) to analyze each generated barcoded BAM file using the default parameters and by increasing to 0.12 the expected error rate for clone identification in order to account for the lower sequencing coverage of the analyzed data^27^. In addition, we provided CHISEL with the available matched-normal germline sample from each patient and phased germline SNPs according to the recommended pipeline by using Eagle2 through the Michigan Imputation Server with the Haplotype Reference Consortium (HRC) reference panel (v.r1.1 2016). CHISEL inferred allele- and haplotype-specific copy numbers per cell and used these results to group cells into distinct tumor clones, while excluding outliers and likely noisy cells. To determine fraction of aberrant genome (genome affected by SCNAs), we defined aberrant as any non-diploid genomic region (i.e., allele-specific copy numbers different than {1, 1}) in tumors not affected by WGDs (NCH-OS-10, NCH-OS-4, and NCH-OS-7) or any non-tetraploid genomic region (i.e., allele-specific copy numbers different than {2, 2}) in tumors affected by WGDs (NCH-OS-8, OS-17, NCH-OS-11, SJOS046149_X2, SJOS003939_X2 and SJOS003939_X1). We defined deletions as previously described in cancer evolutionary studies^26,52–54^. We say that a genomic region in a cell is affected by a deletion when any of the two allele-specific copy numbers inferred by CHISEL is lower than the expected allele-specific copy number (1 for non-WGD tumors or 2 for tumors affected by WGD). Conversely, a genomic region is amplified when any of the two allele-specific copy numbers is higher than expected.

### Reconstruction of copy-number trees

We reconstructed copy-number trees for tumor samples NCH-OS-4 (tibia), NCH-OS-7 (flank) and NCH-OS-7 (tibia), to describe the phylogenetic relationships between distinct tumor clones inferred by CHISEL based on SCNAs using the same procedure proposed in previous studies^27^. Briefly, we reconstructed the trees using the maximum parsimony model of interval events for SCNAs^52,53^ and the copy-number profiles of each inferred clone. These copy number profiles were obtained as the consensus across the inferred haplotype-specific copy numbers derived by CHISEL for all the cells in the same clone, where we also considered the occurrence of WGDs predicted by CHISEL. We classified copy-number events as deletions (i.e., del), as LOH which are deletions resulting in the complete loss of all copies of one allele (loh), as copy-neutral LOH which are LOHs in which the retained allele is simultaneously amplified, and as gains (gain).

### SCNA calling on whole genome data

To compare SCNA patterns across multiple tumor samples from the same patients, we downloaded a total of 47 whole genome sequence datasets from St. Jude’s DNAnexus from 14 patients including germline data and multiple tumor samples (diagnosis, relapse, metastasis and xenograft). We also included the seven scSCNA datasets we generated which had matched germline whole genome data in the St. Jude data and treated these as bulk sequencing data for this analysis. We used samtools^55^ to convert the bam files to fastq and aligned all datasets to a joint hg38/mm10 reference. We filtered out all mouse sequences and removed PCR duplicates. We then called SCNAs with Varscan^56^. Next, we combined all SCNA data by calculating the median copy number for 1,000 bp non-overlapping bins. Correlation between samples was calculated using the cor function in R and the resulting output was plotted as a heatmap using the pheatmap R package (https://github.com/raivokolde/pheatmap).

### SNP calling on whole genome data

To assess genetic heterogeneity of all samples, we produced phylogenetic trees from SNP data. We used the bam alignment files produced during the SCNA calling analysis and called SNPs using bcftools’ mpileup function^55^. We removed SNP calls with a quality below 20 and read depth below 20, and then generated vcf files using bcftools^55^. To check TP53 status we merged the SNP calls with known SNPs from ClinVar^57^ and kept SNPs with a clinical significance (CLNSIG) of “Pathogenic”^58^.

## Supporting information

supplemental figures with legends

## Data and code availability

All the processed data, scripts and results from CHISEL are available on GitHub at https://github.com/kidcancerlab/sc-OsteoCNAs. Whole genome sequencing data for pediatric relapse tumor samples used for analysis in this study were obtained from St. Jude Cloud^38–40^.

## Acknowledgments

We thank our funding sources. This work was generously supported by NIH/NCI K08CA201638 (R.D.R.), R01CA260178 (R.D.R.) and U24CA248453 (B.J.R.), St. Baldrick’s Foundation Scholar Award (R.D.R.), Hyundai Hope on Wheels Young Investigator Award (R.D.R.), Cancer Free Kids Foundation (R.D.R.), Steps for Sarcoma Foundation (R.D.R.), Sarcoma Foundation of America (R.D.R.), a Pelotonia Fellowship (S.R.), a Nationwide Children’s Director’s Strategic Development Fund, and an NIH CTSA Grant UL1TR002733. S.Z. is a Cancer Research UK Career Development Fellow (Award Reference RCCCDF-Nov21\100005) and is also supported by Rosetrees Trust (Grant Reference M917) and by a Cancer Research UK UCL Centre Non-Clinical Training Award (CANTAC721\100022).

## References

1. Casali, P. G. et al. Bone sarcomas: ESMO-PaedCan-EURACAN Clinical Practice Guidelines for diagnosis, treatment and follow-up. Annals of Oncology 29, iv79–iv95 (2018).

2. Bridge, J. A. et al. Cytogenetic findings in 73 osteosarcoma specimens and a review of the literature. Cancer Genet Cytogenet 95, 74–87 (1997).

3. Chen, X. et al. Recurrent somatic structural variations contribute to tumorigenesis in pediatric osteosarcoma. Cell Rep 7, 104–112 (2014).

4. Squire, J. A. et al. High-resolution mapping of amplifications and deletions in pediatric osteosarcoma by use of CGH analysis of cDNA microarrays. Genes Chromosomes Cancer 38, 215–225 (2003).

5. Sayles, L. C. et al. Genome-informed targeted therapy for osteosarcoma. Cancer Discov 9, 46–63 (2019).

6. Stephens, P. J. et al. Massive genomic rearrangement acquired in a single catastrophic event during cancer development. Cell 144, 27–40 (2011).

7. Meyerson, M. & Pellman, D. Cancer genomes evolve by pulverizing single chromosomes. Cell vol. 144 9–10 Preprint at https://doi.org/10.1016/j.cell.2010.12.025 (2011).

8. Li, Y. et al. Constitutional and somatic rearrangement of chromosome 21 in acute lymphoblastic leukaemia. Nature 508, 98–102 (2014).

9. Behjati, S. et al. Recurrent mutation of IGF signalling genes and distinct patterns of genomic rearrangement in osteosarcoma. Nat Commun 8, 1–8 (2017).

10. Perry, J. A. et al. Complementary genomic approaches highlight the PI3K/mTOR pathway as a common vulnerability in osteosarcoma. Proc Natl Acad Sci U S A 111, E5564–E5573 (2014).

11. Zhao, Y. et al. Single-cell RNA sequencing reveals the impact of chromosomal instability on glioblastoma cancer stem cells. BMC Med Genomics 12, 79 (2019).

12. Bakker, B. et al. Single-cell sequencing reveals karyotype heterogeneity in murine and human malignancies. (2016) doi:10.1186/s13059-016-0971-7.

13. Watkins, T. B. K. et al. Pervasive chromosomal instability and karyotype order in tumour evolution. Nature 587, 126–132 (2020).

14. Bach, D. H., Zhang, W. & Sood, A. K. Chromosomal instability in tumor initiation and development. Cancer Research vol. 79 3995–4002 Preprint at https://doi.org/10.1158/0008-5472.CAN-18-3235 (2019).

15. Fearon, E. R. & Vogelstein, B. A genetic model for colorectal tumorigenesis. Cell 61, 759–767 (1990).

16. Hoglund, M. et al. Multivariate analyses of genomic imbalances in solid tumors reveal distinct and converging pathways of karyotypic evolution. Genes Chromosomes Cancer 31, 156–171 (2001).

17. Umbreit, N. T. et al. Mechanisms generating cancer genome complexity from a single cell division error. Science (1979) 368, (2020).

18. Gao, R. et al. Punctuated copy number evolution and clonal stasis in triple-negative breast cancer. Nat Genet 48, 1119–1130 (2016).

19. Zhou, Y. et al. Single-cell RNA landscape of intratumoral heterogeneity and immunosuppressive microenvironment in advanced osteosarcoma. Nat Commun 11, 1–17 (2020).

20. Liu, Y. et al. Single-Cell Transcriptomics Reveals the Complexity of the Tumor Microenvironment of Treatment-Naive Osteosarcoma. Front Oncol 0, 2818 (2021).

21. Xu, H. et al. Genetic and clonal dissection of osteosarcoma progression and lung metastasis. Int J Cancer 143, 1134–1142 (2018).

22. Negri, G. L. et al. Integrative genomic analysis of matched primary and metastatic pediatric osteosarcoma. J Pathol 249, 319–331 (2019).

23. Wang, Y. & Navin, N. E. Advances and Applications of Single Cell Sequencing Technologies. Mol Cell 58, 598 (2015).

24. Laks, E. et al. Clonal Decomposition and DNA Replication States Defined by Scaled Single-Cell Genome Sequencing. Cell 179, 1207–1221.e22 (2019).

25. Tarabichi, M. et al. A practical guide to cancer subclonal reconstruction from DNA sequencing. Nature Methods 2021 18:2 18, 144–155 (2021).

26. Zaccaria, S. & Raphael, B. J. Accurate quantification of copy-number aberrations and whole-genome duplications in multi-sample tumor sequencing data. Nature Communications 2020 11:1 11, 1–13 (2020).

27. Zaccaria, S. & Raphael, B. J. Characterizing allele- and haplotype-specific copy numbers in single cells with CHISEL. Nat Biotechnol (2020) doi:10.1038/s41587-020-0661-6.

28. Andor, N. et al. Joint single cell DNA-seq and RNA-seq of gastric cancer cell lines reveals rules of in vitro evolution. NAR Genom Bioinform 2, (2020).

29. Houghton, P. J. et al. The pediatric preclinical testing program: Description of models and early testing results. Pediatr Blood Cancer (2007) doi:10.1002/pbc.21078.

30. Woo, X. Y. et al. Conservation of copy number profiles during engraftment and passaging of patient-derived cancer xenografts. Nature Genetics 2021 53:1 53, 86–99 (2021).

31. Martin, J. W., Squire, J. A. & Zielenska, M. The genetics of osteosarcoma. Sarcoma 2012, 11 (2012).

32. Minussi, D. C. et al. Breast tumours maintain a reservoir of subclonal diversity during expansion. Nature 592, 302–308 (2021).

33. Zack, T. I. et al. Pan-cancer patterns of somatic copy number alteration. Nat Genet 45, 1134–1140 (2013).

34. López, S. et al. Interplay between whole-genome doubling and the accumulation of deleterious alterations in cancer evolution. Nat Genet 52, 283–293 (2020).

35. Bielski, C. M. et al. Genome doubling shapes the evolution and prognosis of advanced cancers. Nat Genet 50, 1189–1195 (2018).

36. Passerini, V. et al. The presence of extra chromosomes leads to genomic instability. Nat Commun 7, 1–12 (2016).

37. Sheltzer, J. M. A transcriptional and metabolic signature of primary aneuploidy is present in chromosomally unstable cancer cells and informs clinical prognosis. Cancer Res 73, 6401–6412 (2013).

38. McLeod, C. et al. St. Jude Cloud: A Pediatric Cancer Genomic Data-Sharing Ecosystem. Cancer Discov 11, 1082–1099 (2021).

39. Downing, J. R. et al. The Pediatric Cancer Genome Project. Nat Genet 44, 619–622 (2012).

40. Stewart, E. et al. Orthotopic patient-derived xenografts of paediatric solid tumours. Nature 549, 96–100 (2017).

41. Zaccaria, S. & Raphael, B. J. Accurate quantification of copy-number aberrations and whole-genome duplications in multi-sample tumor sequencing data. Nature Communications 2020 11:1 11, 1–13 (2020).

42. Wang, D. et al. Multiregion sequencing reveals the genetic heterogeneity and evolutionary history of osteosarcoma and matched pulmonary metastases. Cancer Res 79, 7–20 (2019).

43. Brady, S. W. et al. The Clonal Evolution of Metastatic Osteosarcoma as Shaped by Cisplatin Treatment. Mol Cancer Res 17, 895–906 (2019).

44. Navin, N. et al. Tumour evolution inferred by single-cell sequencing. Nature 472, 90–95 (2011).

45. Wang, Y. et al. Clonal evolution in breast cancer revealed by single nucleus genome sequencing. Nature 512, 155–160 (2014).

46. Navin, N. E. The first five years of single-cell cancer genomics and beyond. Genome Research vol. 25 1499–1507 Preprint at https://doi.org/10.1101/gr.191098.115 (2015).

47. Gawad, C., Koh, W. & Quake, S. R. Single-cell genome sequencing: current state of the science. Nature Reviews Genetics 2016 17:3 17, 175–188 (2016).

48. Gröbner, S. N. et al. The landscape of genomic alterations across childhood cancers. Nature 555, 321–327 (2018).

49. Ben-David, U. et al. Patient-derived xenografts undergo mouse-specific tumor evolution. Nature Genetics 2017 49:11 49, 1567–1575 (2017).

50. Ma, X. et al. Pan-cancer genome and transcriptome analyses of 1,699 paediatric leukaemias and solid tumours. Nature 555, 371–376 (2018).

51. Rajan, S. et al. Osteosarcoma tumors maintain intratumoral heterogeneity, even while adapting to environmental pressures that drive clonal selection. bioRxiv 2020.11.03.367342 Preprint at https://doi.org/10.1101/2020.11.03.367342 (2020).

52. Schwarz, R. F. et al. Phylogenetic Quantification of Intra-tumour Heterogeneity. PLoS Comput Biol 10, e1003535 (2014).

53. El-Kebir, M. et al. Complexity and algorithms for copy-number evolution problems. Algorithms for Molecular Biology 2017 12:1 12, 1–11 (2017).

54. Zaccaria Simone, El-KebirMohammed, W., K. & J., R. Phylogenetic Copy-Number Factorization of Multiple Tumor Samples. https://home.liebertpub.com/cmb 25, 689–708 (2018).

55. Danecek, P. et al. Twelve years of SAMtools and BCFtools. Gigascience 10, 1–4 (2021).

56. Koboldt, D. C. et al. VarScan 2: Somatic mutation and copy number alteration discovery in cancer by exome sequencing. Genome Res 22, 568–576 (2012).

57. Landrum, M. J. et al. ClinVar: improving access to variant interpretations and supporting evidence. Nucleic Acids Res 46, D1062–D1067 (2018).

58. Tamura, K., Stecher, G. & Kumar, S. MEGA11: Molecular Evolutionary Genetics Analysis Version 11. Mol Biol Evol 38, 3022–3027 (2021).

