## supplemental figures with legends for "Structurally complex osteosarcoma genomes exhibit limited heterogeneity within individual tumors and across evolutionary time"

**Supplemental Figure S1: Cell count and fraction of clones**

A. Quantification of cells captured for single cell DNA sequencing in each of the samples. B. Stacked barplots showing the composition of each sample by clone as determined by CHISEL.

**Supplemental Figure S2: CHISEL output plots**

A-J. Raw, unfiltered copy number profiles (without filtering noisy cells) for the same data shown in Figure 1.

**Supplemental Figure S3: Ploidy plots**

Cell-by-cell determination of ploidy for scDNA sequencing shown in Figure 1.

**Supplemental Figure S4: Proportion of identically assigned SCNAs between samples**

Heatmap showing the proportion of genome showing identical copy numbers between samples using the genomic windows that were called by CHISEL. This excludes windows where both samples were copy number neutral to avoid artificial inflation of values in samples with few SCNAs. Note that these results are, by nature, a very conservative quantification of the minimal degree of pairwise shared identity due to the inherent noise and variability within the data (and should be considered an "at least" value) but are useful for comparing similarity among groups of samples.

**Supplemental Figure S5: Fraction of altered genome density plots**

A-E. Ridgeplots show the fraction of the genome that is (A) aberrant, (B) has subclonal variation, exhibits (C) amplification or (D) deletion, and the (E) ploidy of each sample included in the scDNA sequencing dataset.

**Supplemental Figure S6: TP53 copy number in single-cell data**

Copy number status determination for chromosome 17 shows widespread deletion and LOH at the p53 locus cell-by-cell from the scDNA data.

**Supplemental Figure S7: CHISEL plot for SJOS003939 samples**

High-resolution copy number determinations for CHISEL analysis of the pooled SJOS003939 sample set.

**Supplemental Figure S8: St. Jude bulk data genome SCNA plots by patient**

A-J. Copy number plots from patients from the St. Jude dataset where WGS data was available from biopsy/resection and/or PDX samples separated by space (metastasis) and/or time. Inset numbers show estimations of tumor purity and SCNA-specific clone composition determined by HATCHET where available.

Supplemental Figure 1

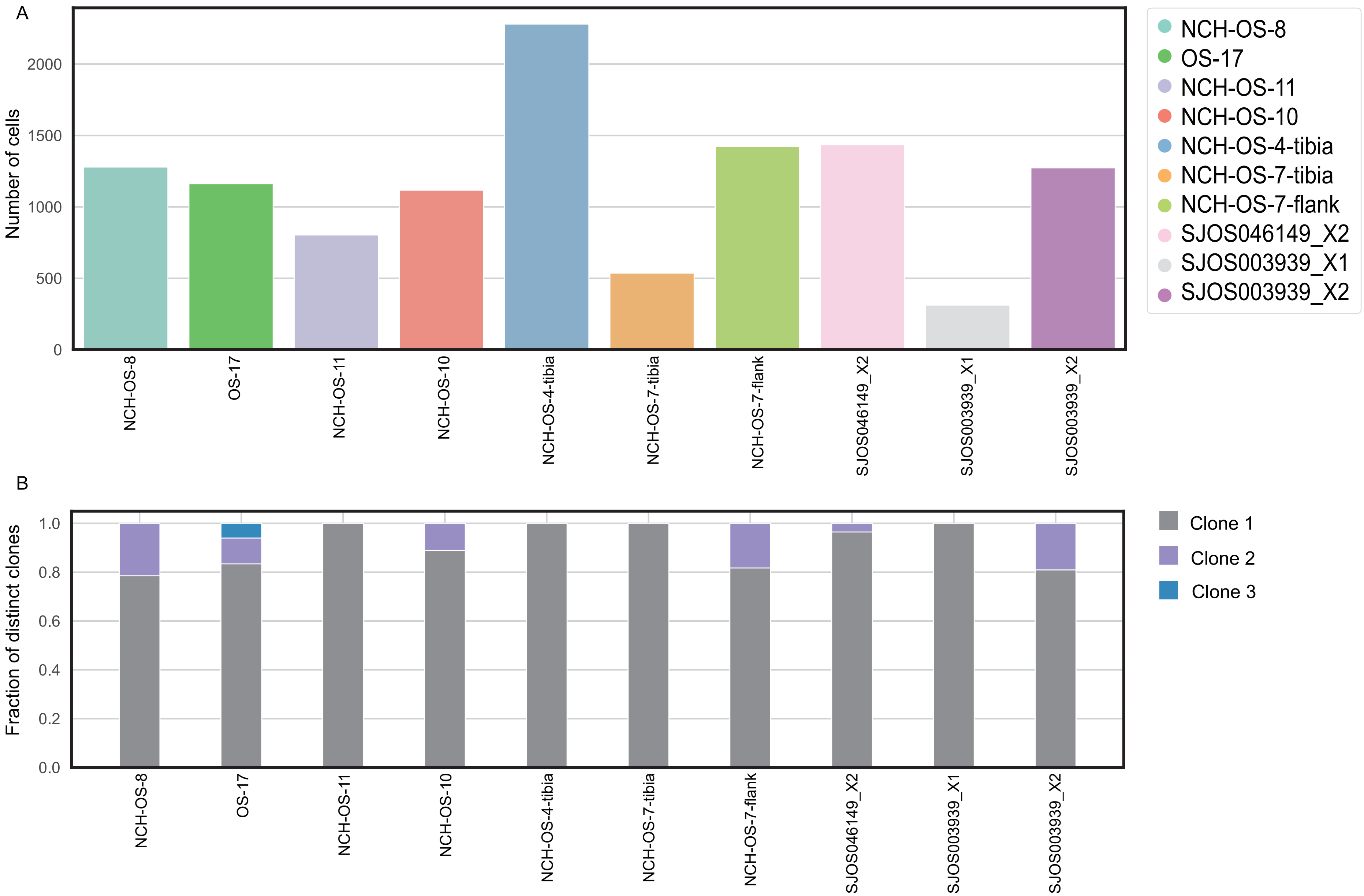

Supplemental Figure 2

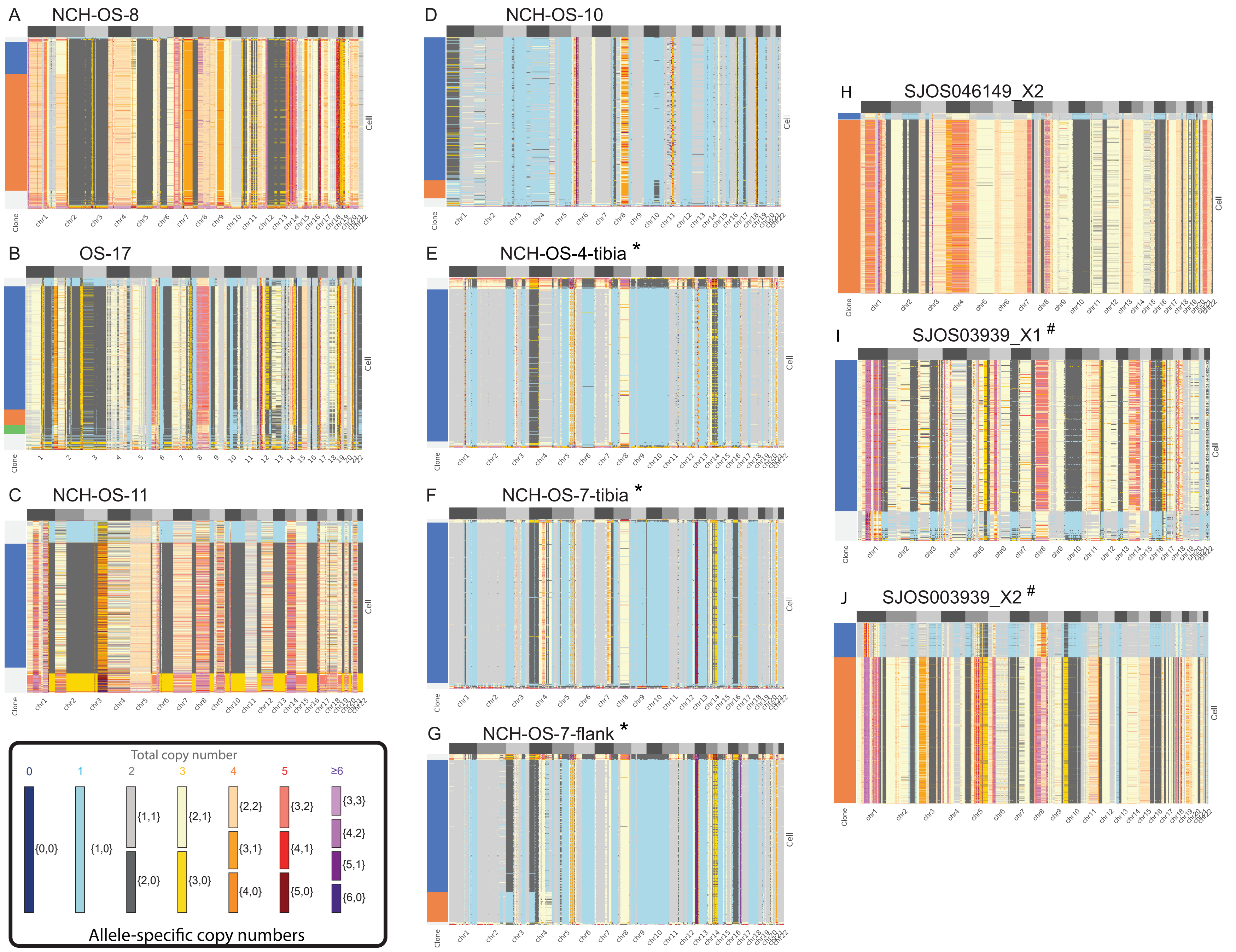

### Supplemental Figure 3

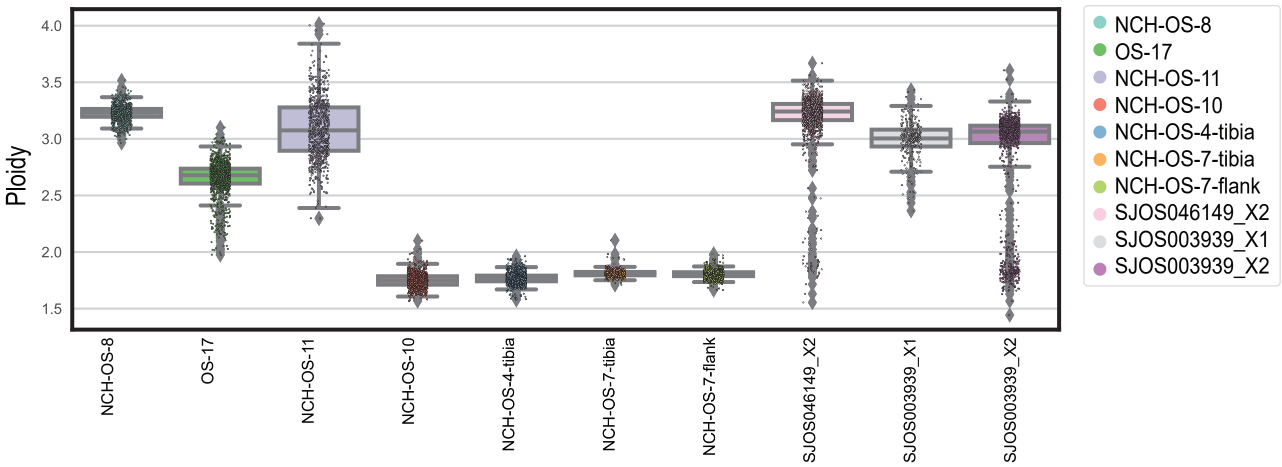

Supplemental Figure 4

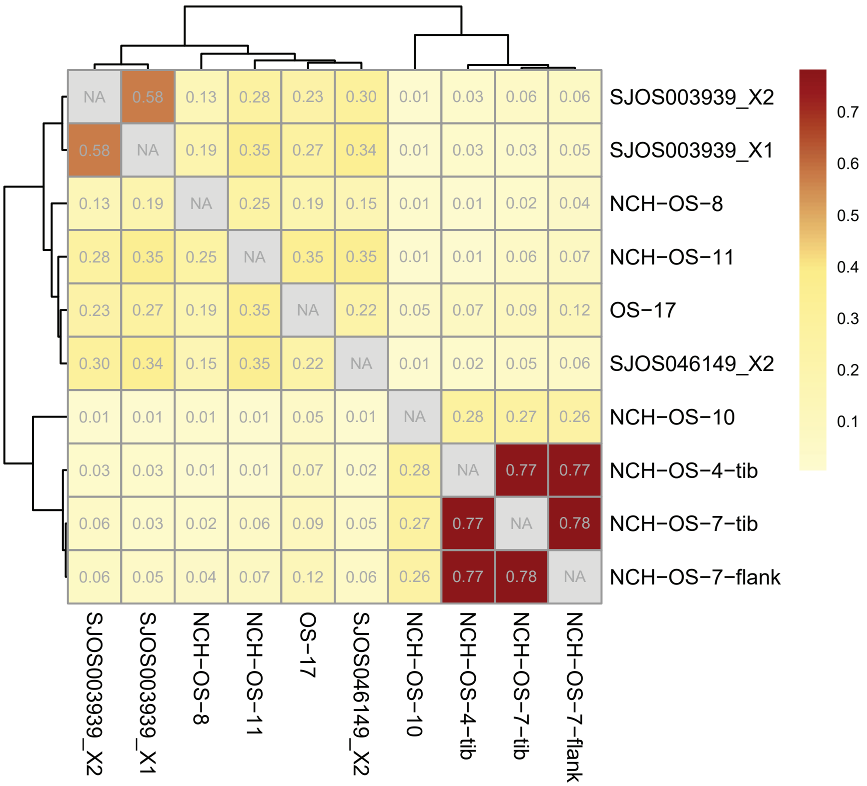

Supplemental Figure 5

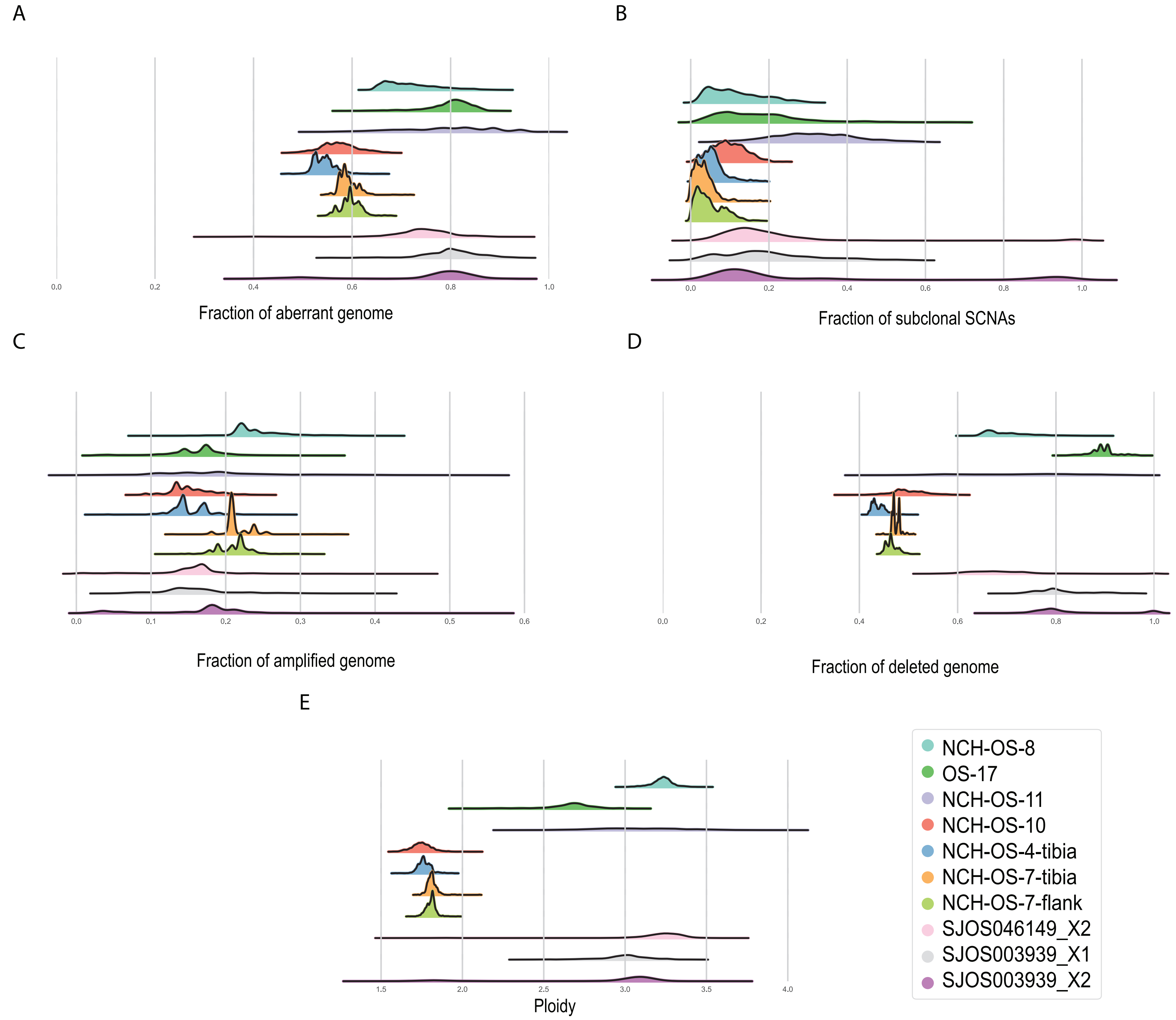

Supplemental Figure 6

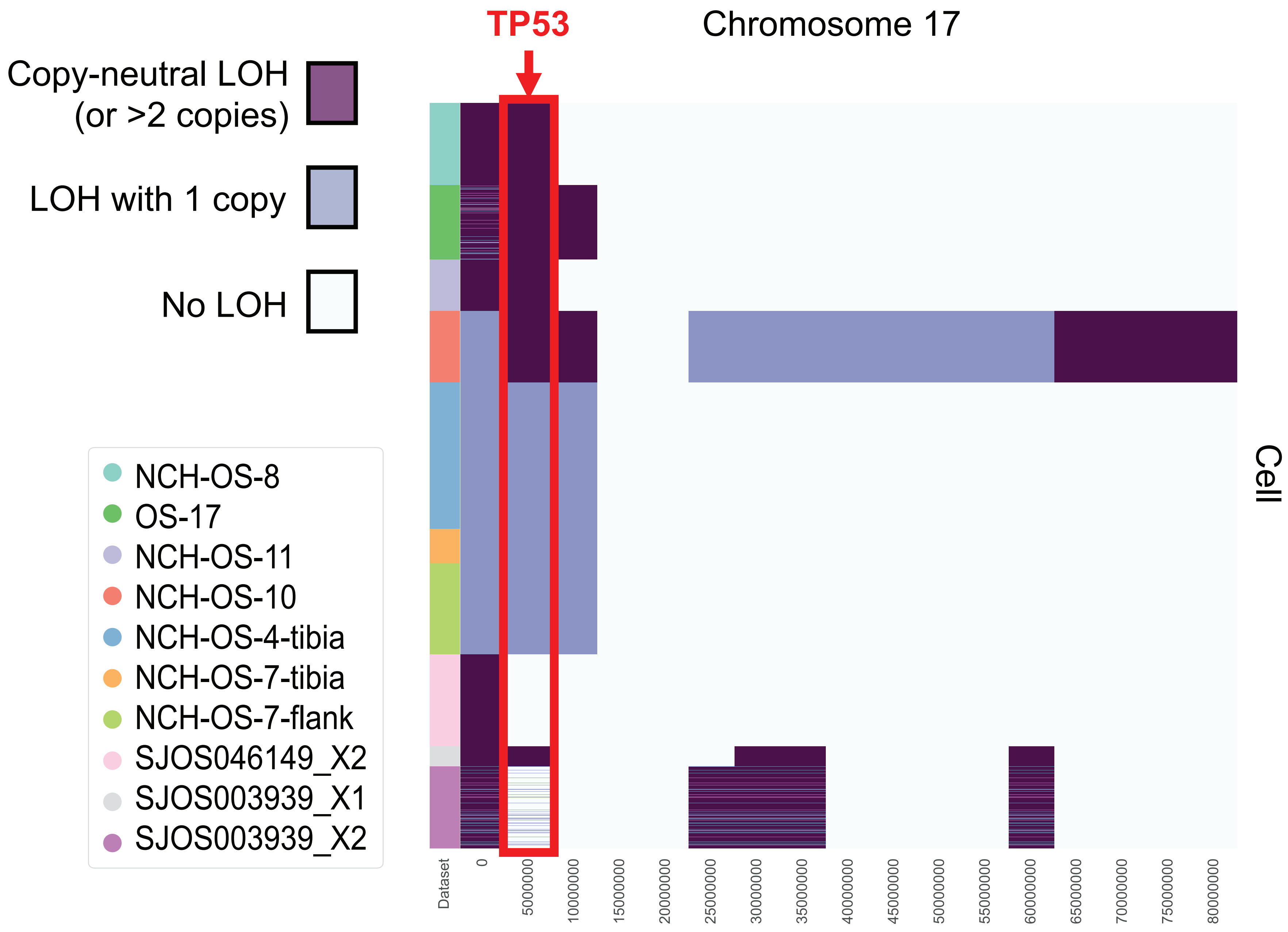

Supplemental Figure 7

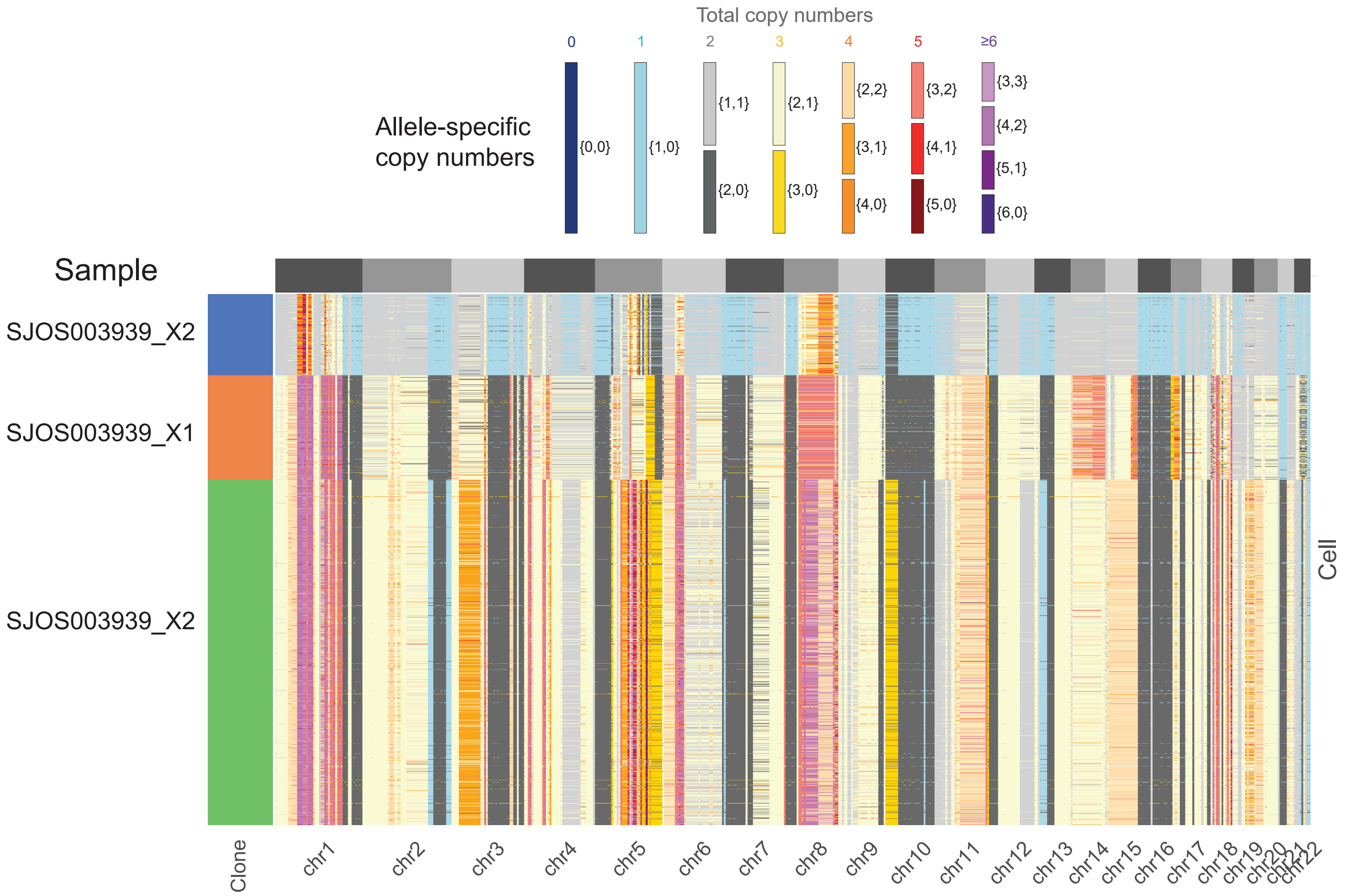

Supplemental Figure 8A

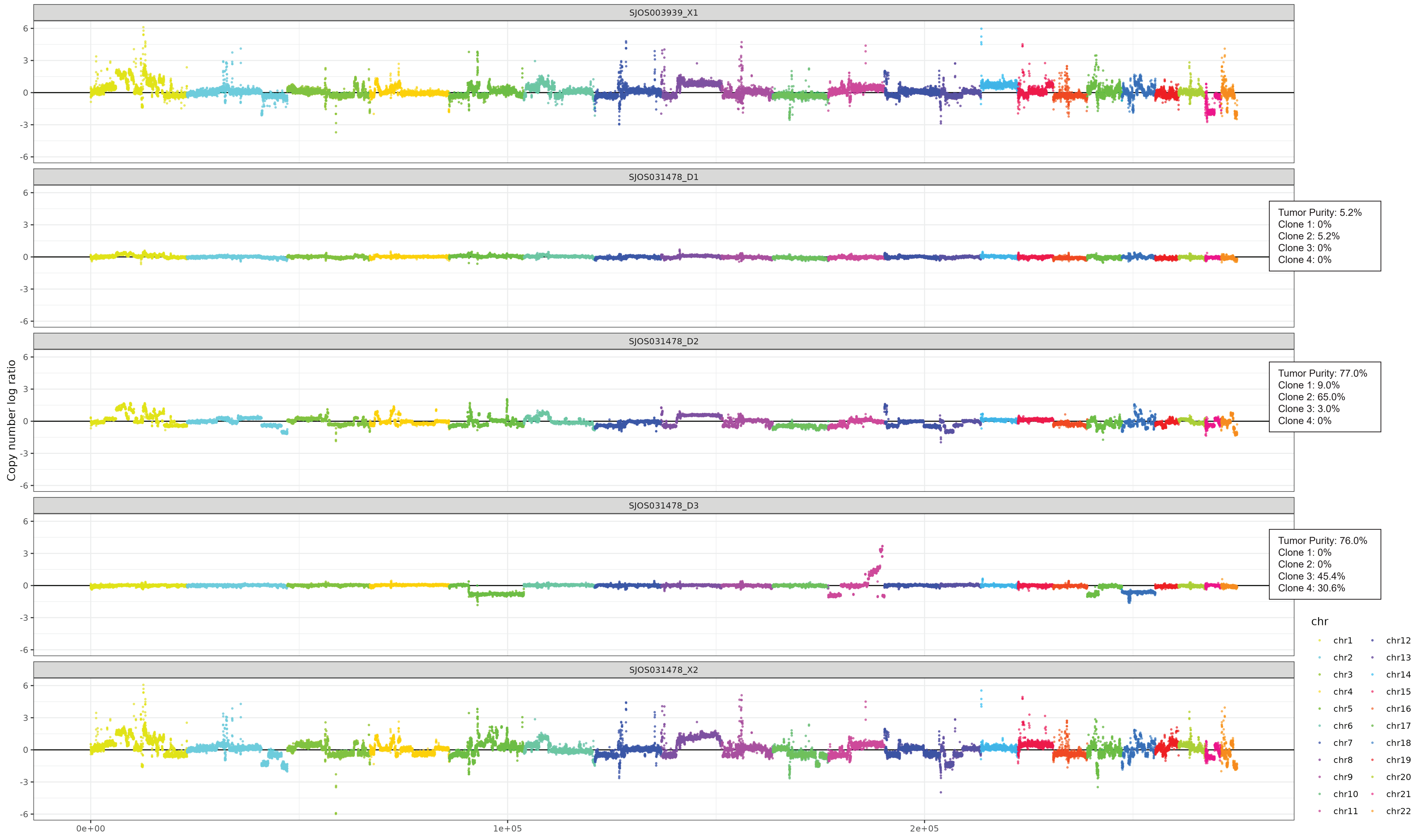

Supplemental Figure 8B

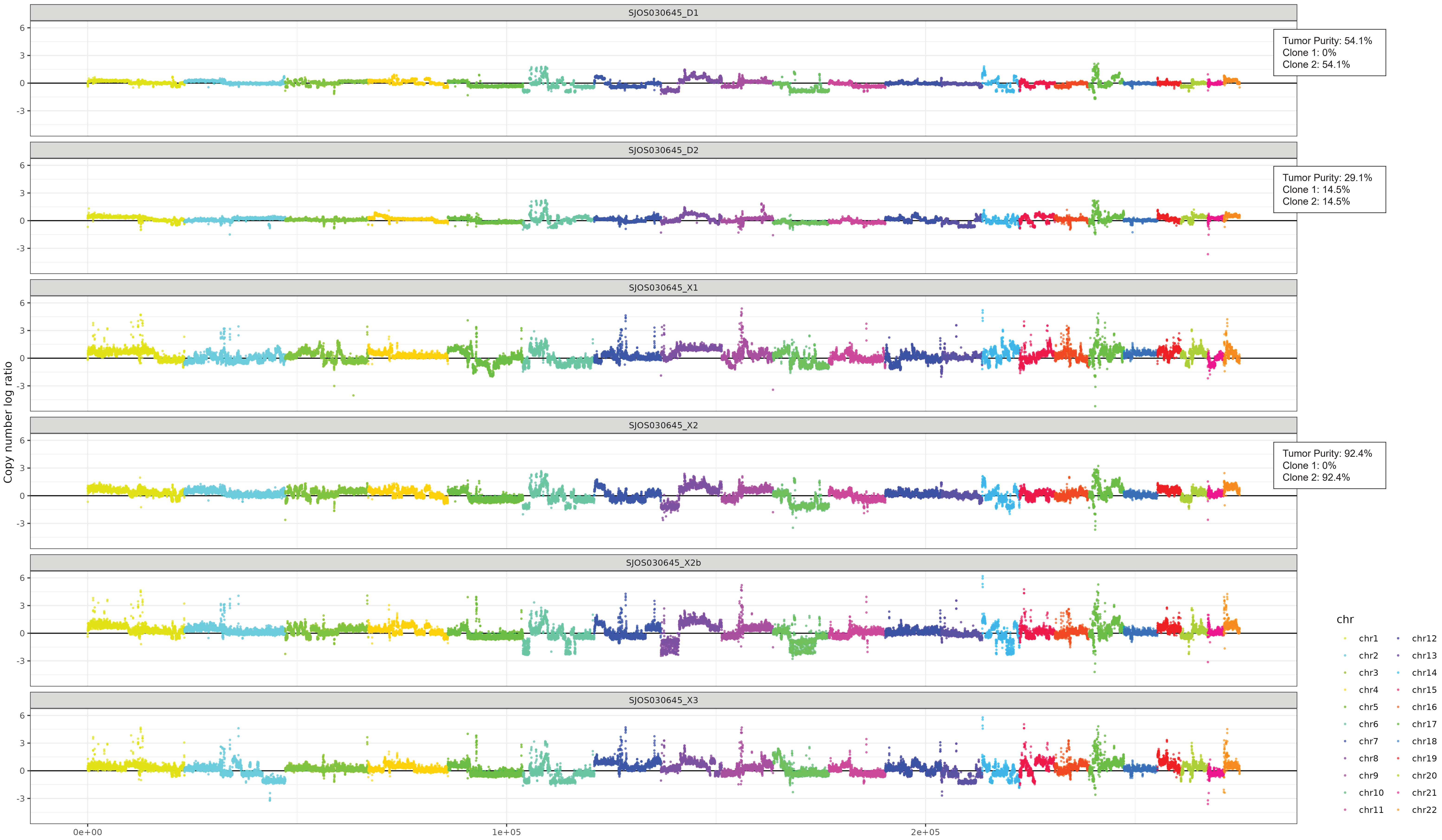

Supplemental Figure 8C

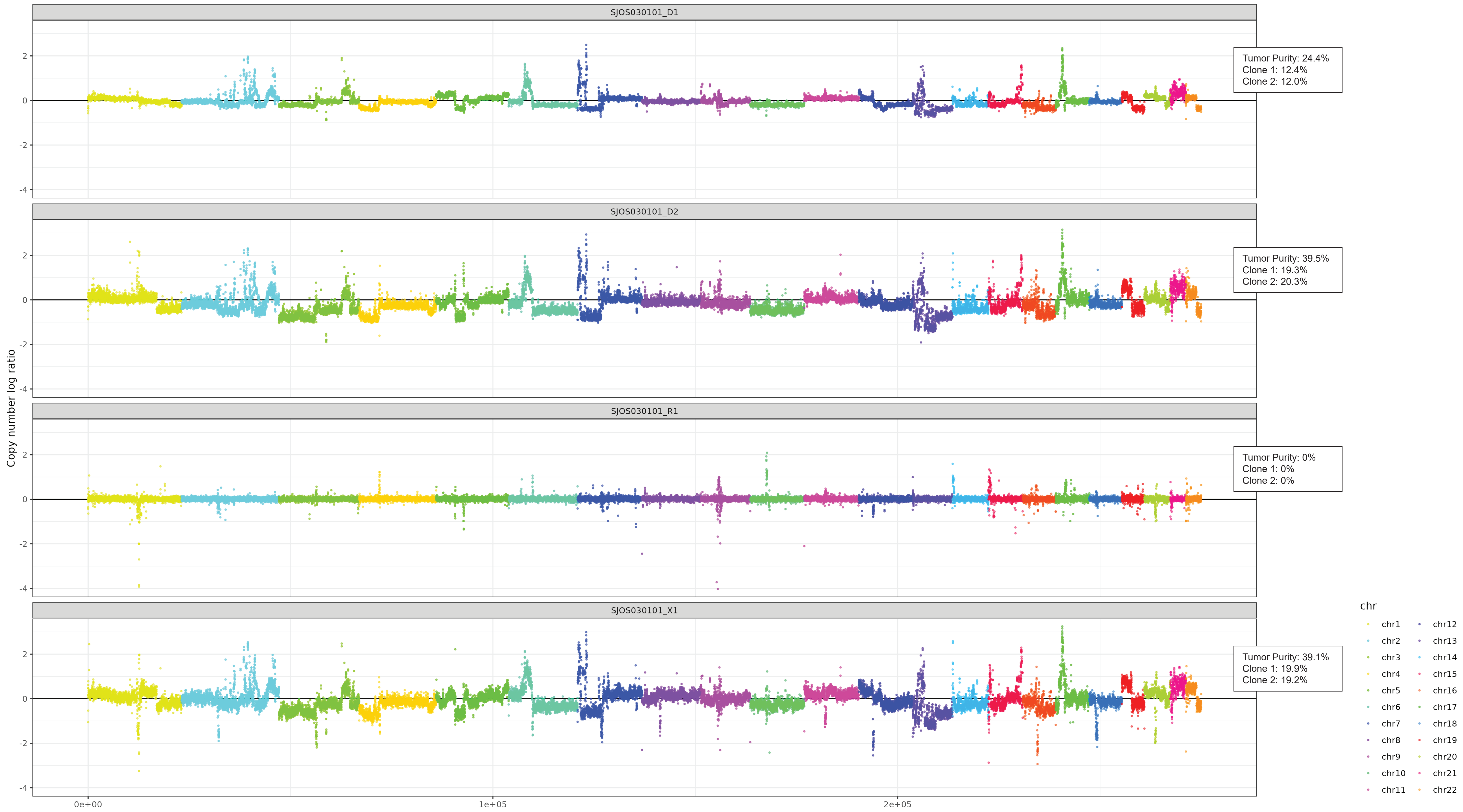

Supplemental Figure 8D

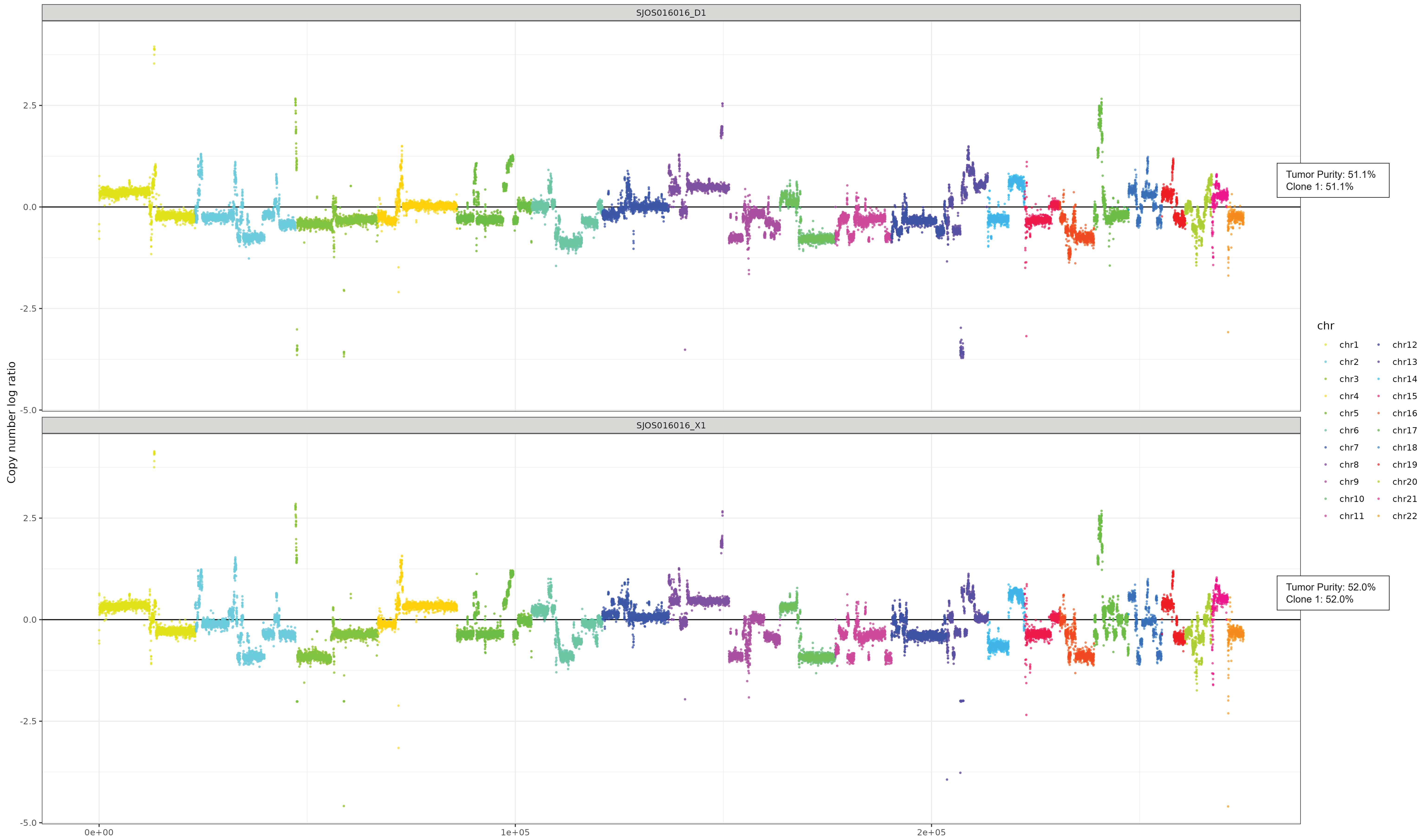

Supplemental Figure 8E

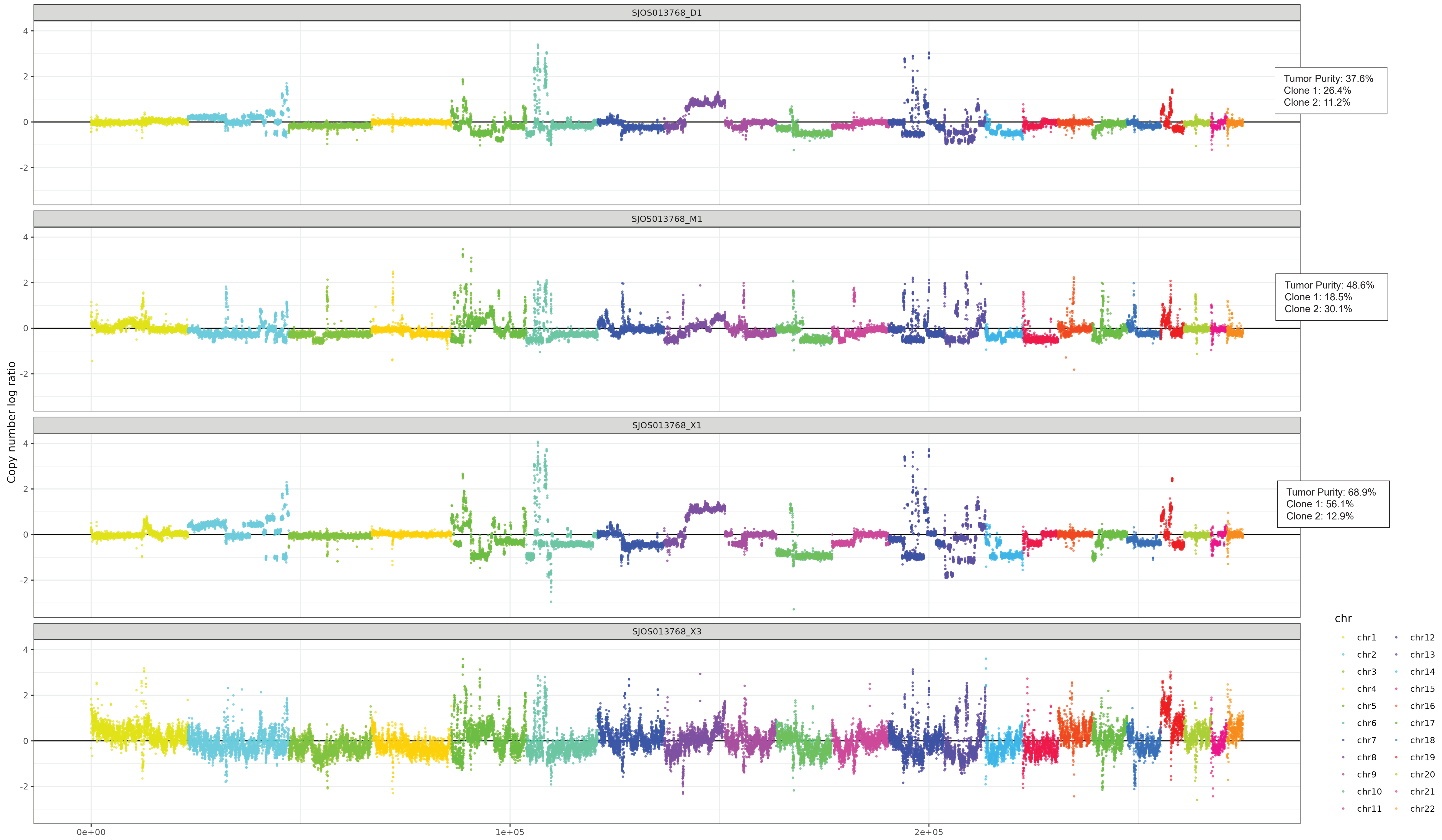

Supplemental Figure 8F

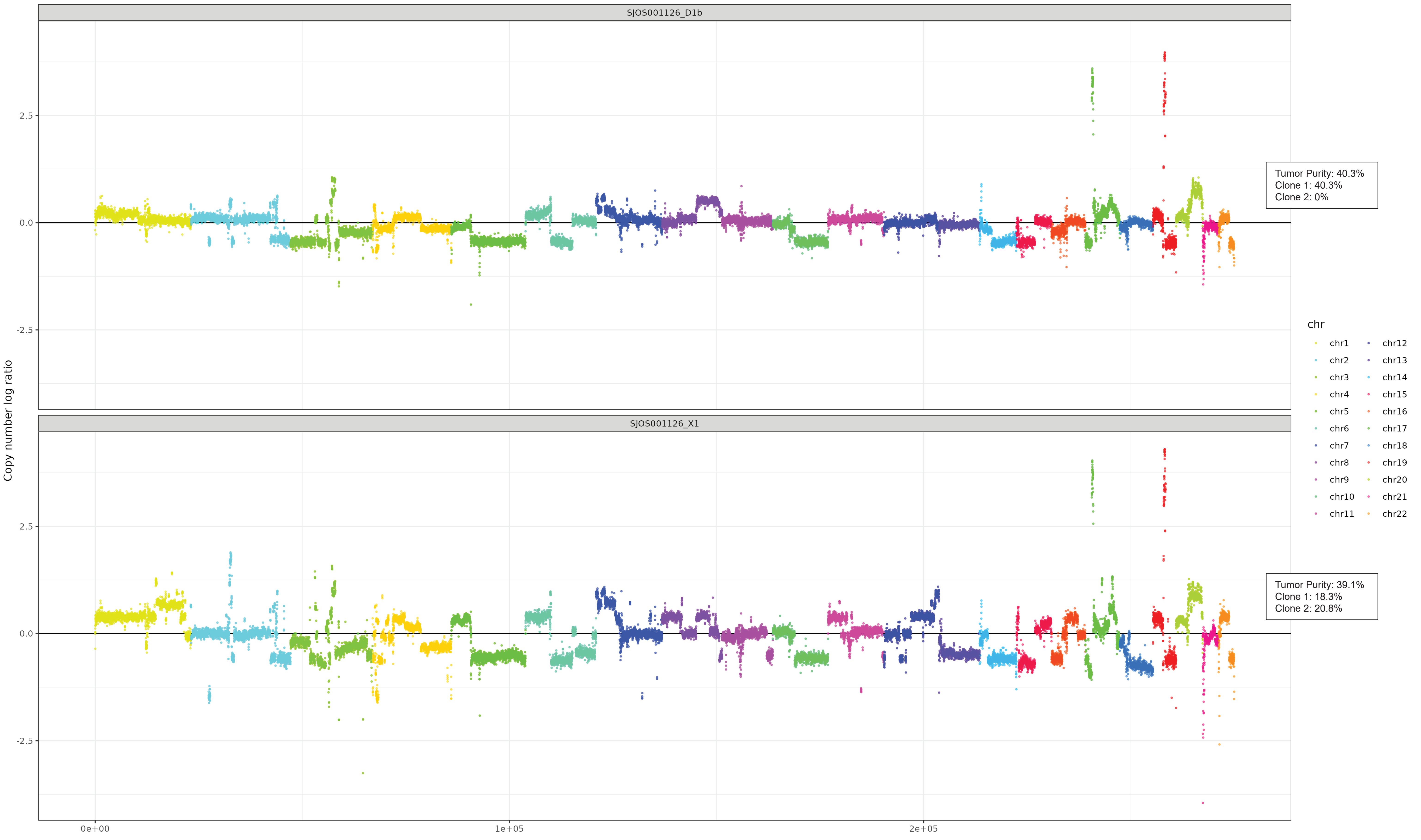

Supplemental Figure 8G

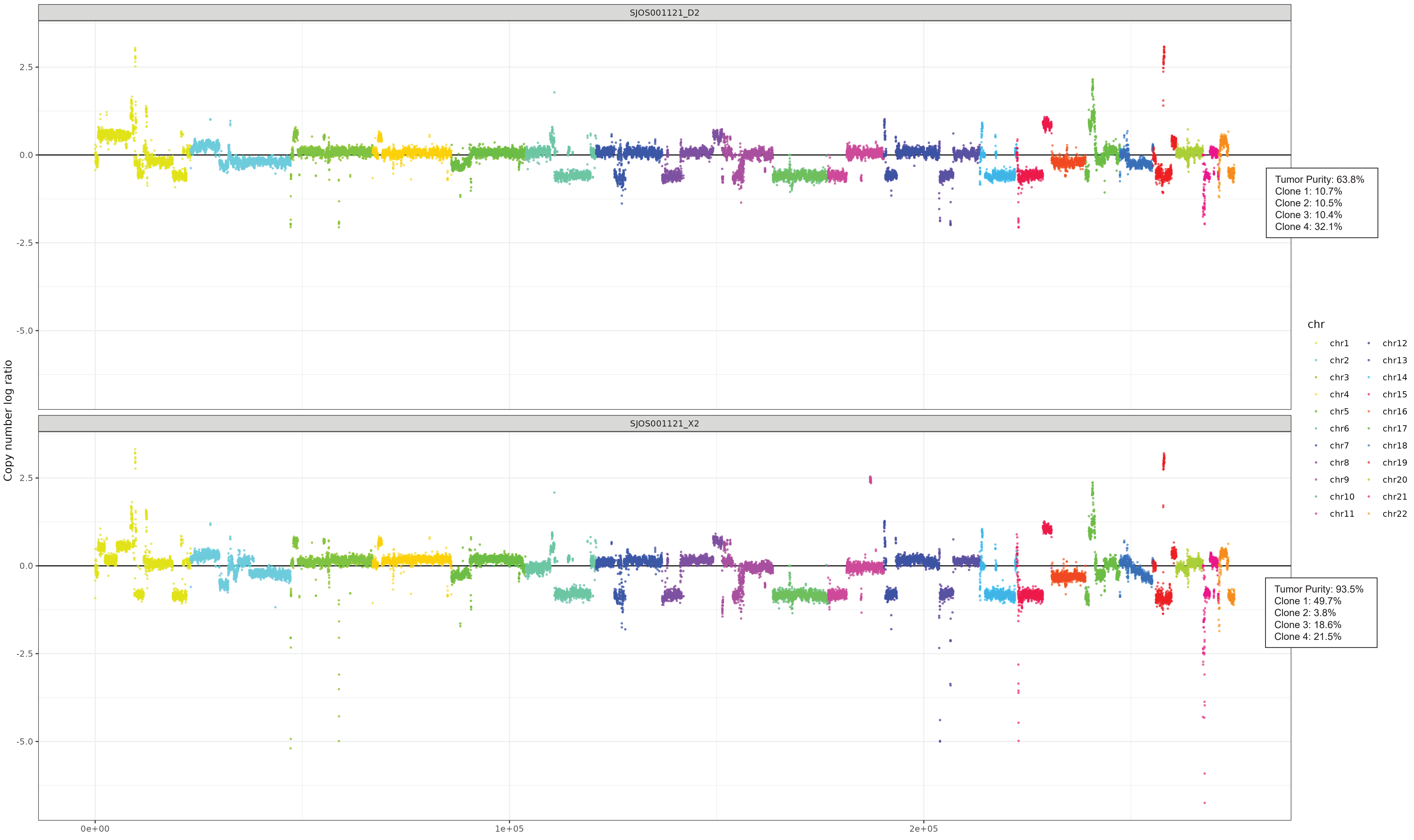

Supplemental Figure 8H

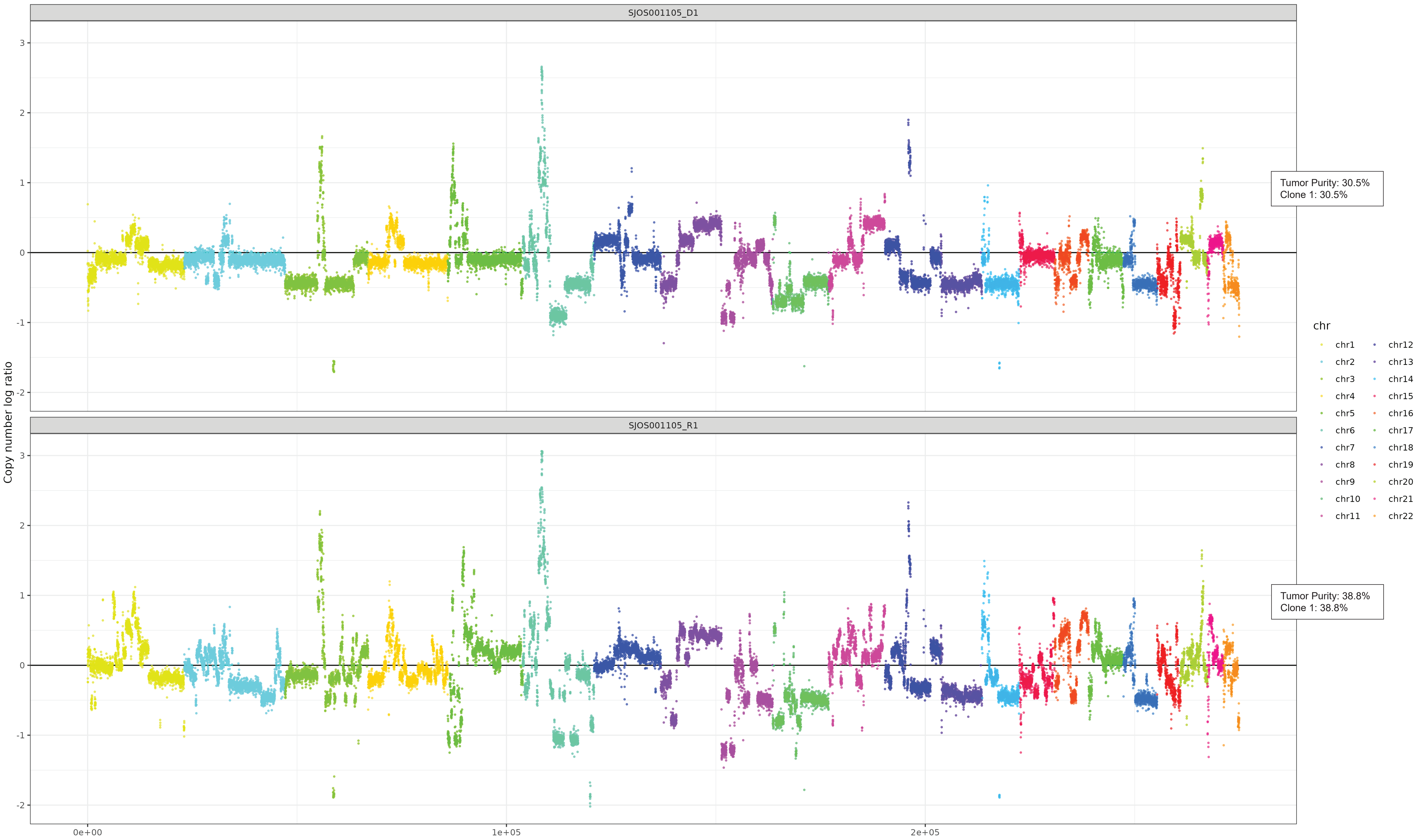

Supplemental Figure 8I

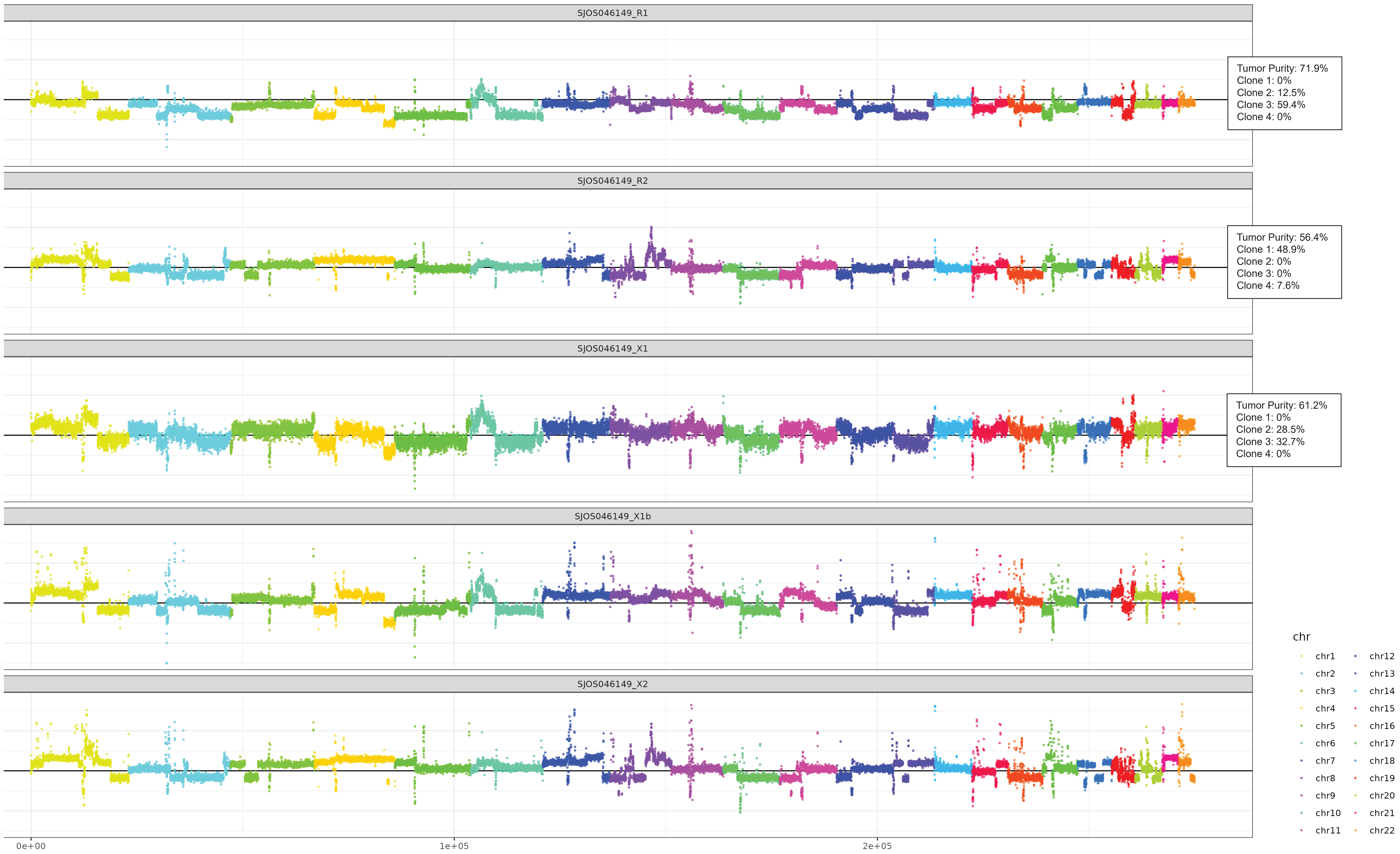

Supplemental Figure 8J

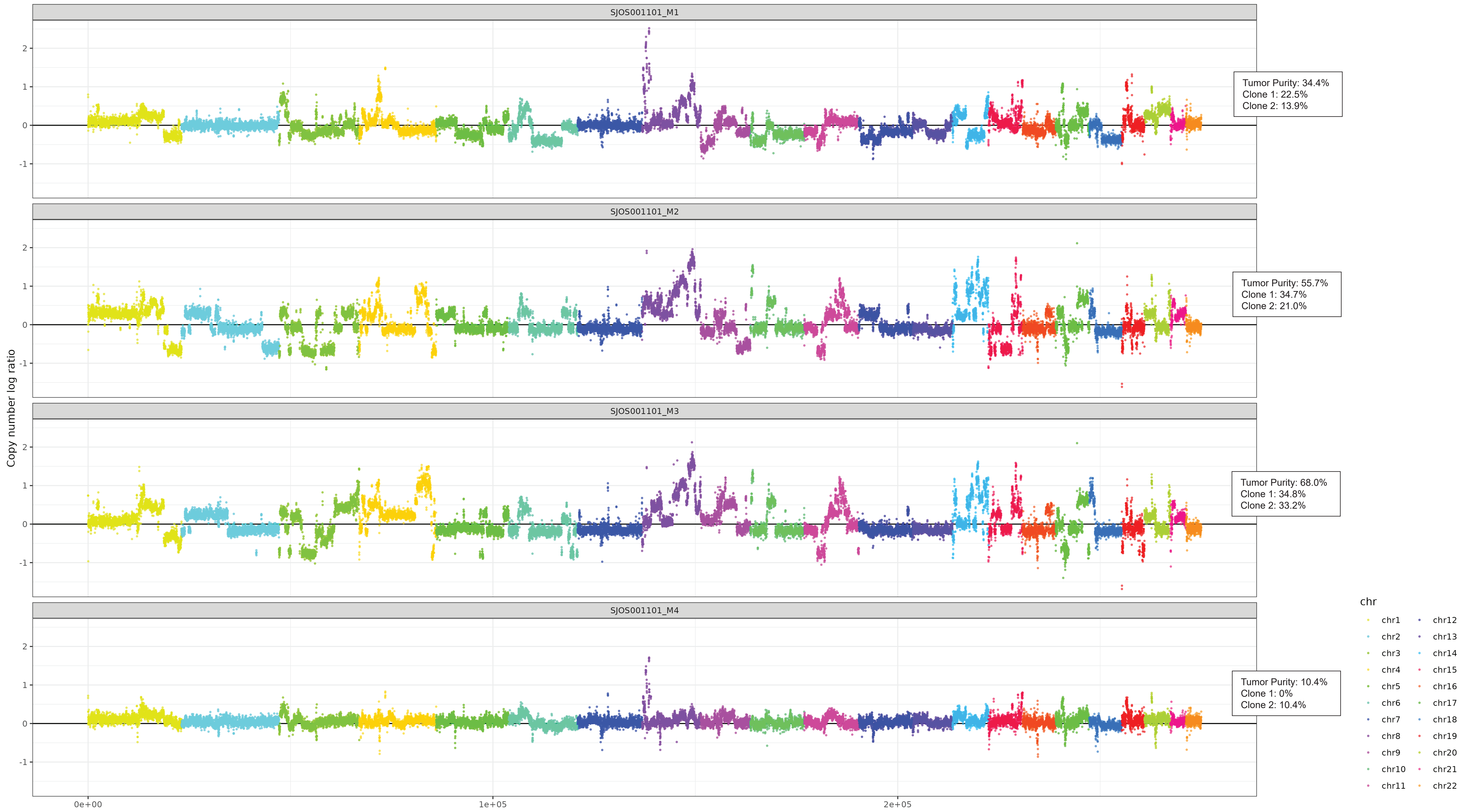

Supplemental Figure 8A

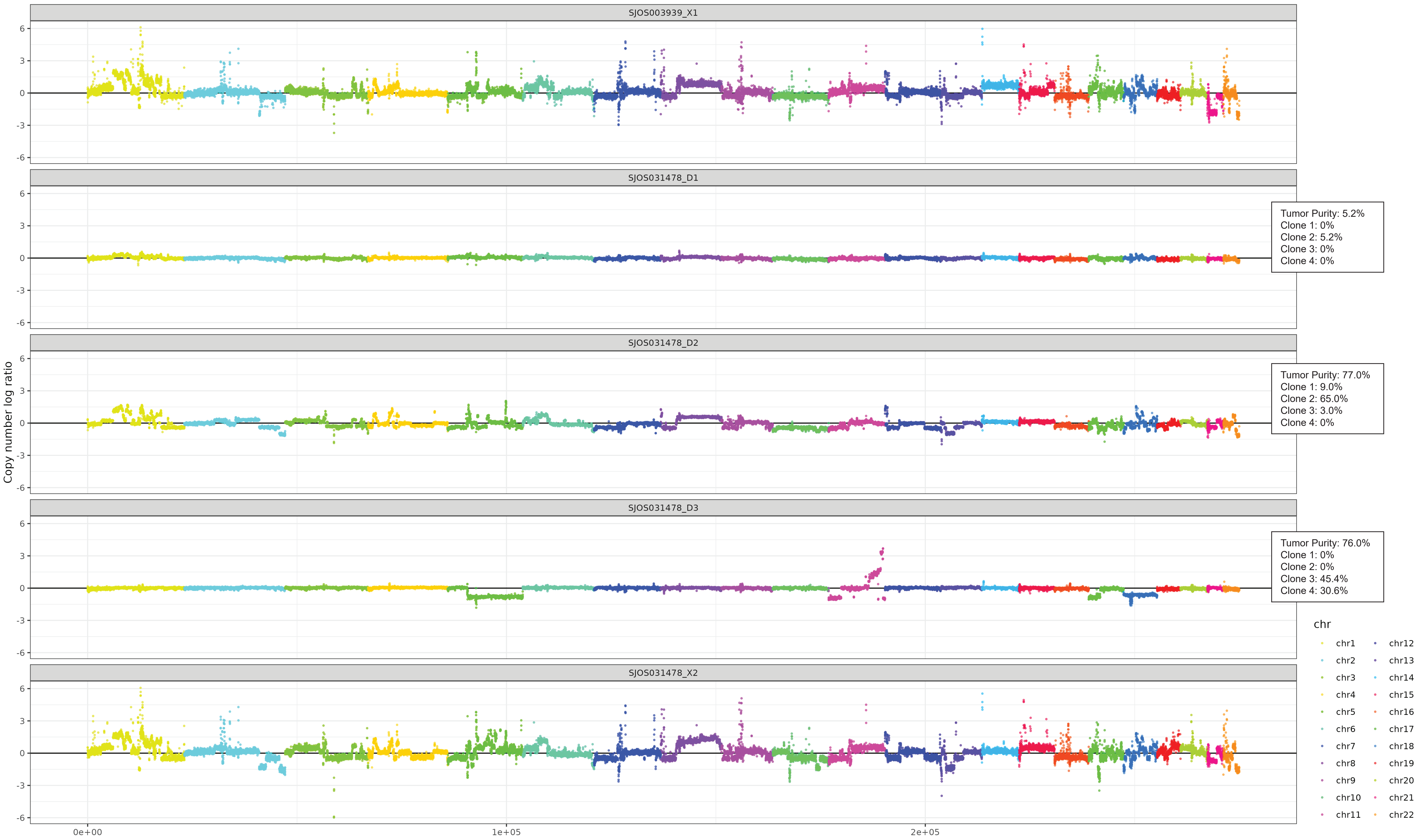
